# Cell wall carbohydrate dynamics during the differentiation of infection structures by the apple scab fungus, *Venturia inaequalis*

**DOI:** 10.1101/2022.09.21.508768

**Authors:** Mercedes Rocafort, Vaibhav Srivastava, Joanna K. Bowen, Sara M. Díaz-Moreno, Vincent Bulone, Kim M. Plummer, Paul W. Sutherland, Marilyn A. Anderson, Rosie E. Bradshaw, Carl H. Mesarich

**Author notes:** Corresponding author: Carl H. Mesarich. Postal address: School of Agriculture and Environment, Massey University, Private Bag 11222, Palmerston North 4442, New Zealand. College of Medicine and Public Health, Flinders University, Bedford Park, South Australia 5042, Australia (Vincent Bulone).

## Abstract

Scab, caused by the biotrophic fungal pathogen *Venturia inaequalis*, is the most economically important disease of apples. During infection, *V. inaequalis* colonizes the subcuticular host environment, where it develops specialized infection structures called runner hyphae and stromata. These structures are thought to be involved in nutrient acquisition and effector (virulence factor) delivery, but also give rise to conidia that further the infection cycle. Despite their importance, very little is known about how these structures are differentiated. Likewise, nothing is known about how these structures are protected from host defences or recognition by the host immune system. To better understand these processes, we first performed a glycosidic linkage analysis of sporulating tubular hyphae from *V. inaequalis* developed in culture. This analysis revealed that the *V. inaequalis* cell wall is mostly composed of glucans (44%) and mannans (37%), whereas chitin represents a much smaller proportion (4%). Next, we used transcriptomics and confocal laser scanning microscopy to provide insights into the cell wall carbohydrate composition of runner hyphae and stromata. These analyses revealed that, during subcuticular host colonization, genes of *V. inaequalis* putatively associated with the biosynthesis of immunogenic carbohydrates, such as chitin and β-1,6-glucan, are down-regulated relative to growth in culture, while on the surface of runner hyphae and stromata, chitin is deacetylated to the less immunogenic carbohydrate, chitosan. These changes are anticipated to enable the subcuticular differentiation of runner hyphae and stromata by *V. inaequalis*, as well as to protect these structures from host defences and recognition by the host immune system.

**Importance:** Plant-pathogenic fungi are a major threat to food security. Among these are subcuticular pathogens, which often cause latent asymptomatic infections, making them difficult to control. A key feature of these pathogens is their ability to differentiate specialized subcuticular infection structures that, to date, remain largely understudied. This is typified by *Venturia inaequalis*, which causes scab, the most economically important disease of apples. In this study, we show that, during subcuticular host colonization, *V. inaequalis* down-regulates genes associated with the biosynthesis of two immunogenic cell wall carbohydrates, chitin and β-1,6-glucan, and coats its infection structures with a less-immunogenic carbohydrate, chitosan. These changes are anticipated to enable subcuticular host colonization by *V. inaequalis* and provide a foundation for understanding subcuticular host colonization by other plant-pathogenic fungi. Such an understanding is important, as it may inform the development of novel control strategies against subcuticular plant-pathogenic fungi.

## Introduction

Scab, caused by the fungus *Venturia inaequalis*, is one of the most devastating diseases of apples worldwide (1, 2). Under favourable conditions, disease symptoms emerge as brown-green lesions on leaves, buds and fruit, rendering the fruit unmarketable and reducing crop yield by up to 70% (3, 4). Scab is also the most expensive disease of apples to control, with up to 20 fungicide treatments required each year (4, 5). This intensive fungicide use has accelerated the development of fungicide resistance in *V. inaequalis* and has increased production costs for growers (6). While some disease-resistant apple cultivars have been developed, their use has been limited due to the rapid emergence of resistance-breaking strains of *V. inaequalis* in the field (7). Thus, there is an urgent need to develop durable control strategies against scab.

*V. inaequalis* is a biotrophic pathogen that colonizes the subcuticular (apoplastic) host environment located between the cuticle and underlying epidermal cells of apple tissues (1, 4). During colonization, *V. inaequalis* develops specialized subcuticular infection structures called runner hyphae and stromata (8-10). These structures are non-melanized and are different from regular tubular hyphae developed on the host surface (9). Indeed, runner hyphae are wider and flatter than regular tubular hyphae and are often fused along their length to form ‘hyphal superhighways’ (9), while stromata are multi-layered pseudoparenchymatous structures that are the result of a switch from polar tip extension to non-polar lateral division (9). In terms of functionality, stromata give rise to asexual conidia that further the infection cycle but are also thought to be involved in nutrient acquisition and effector (virulence factor) delivery. Runner hyphae, on the other hand, enable the fungus to radiate out from the initial site of host penetration, acting as a base from which additional stromata can be differentiated (9).

Notably, subcuticular infection structures are also produced by other crop-infecting members of the *Venturia* genus (11-14), as well as several other species of plant-pathogenic fungi. The latter include *Diplocarpon rosae*, which causes black spot disease of roses (15), and *Rhynchosporium secalis*, which causes barley and rye scald (16, 17). Despite these observations, very little is known about how these structures are differentiated or how the fungal cell wall is remodelled during this process. Strikingly, *V. inaequalis* can develop infection-like structures inside cellophane membranes (CMs) that are reminiscent of those formed *in planta* (9). This contrasts with growth on the CM surface, where the fungus develops tubular hyphae like those formed on the surface of apple tissues (9). This finding suggests that CMs can be used as an in culture model for studying the differentiation of subcuticular infection structures and the dynamics of cell wall remodelling.

The fungal cell wall is an external barrier that plays an essential role in fungal growth and morphogenesis (18, 19). For plant-pathogenic fungi, the cell wall also has an important role in protection, as it is the first structure to encounter the hostile apoplastic environment of the host (20). Despite this importance, its composition and biosynthesis are still poorly understood, especially for non-model filamentous pathogens (19). While the structure and composition of the fungal cell wall differs between species, it is typically comprised of a polysaccharide and protein matrix, with glucans, chitin and mannans the main components (18). Crucially, some of these carbohydrates, such as chitin and β-glucan, are strong elicitors of the plant immune system, with defence responses initiated upon their recognition as microbe-associated molecular patterns (MAMPs) by cell surface-localized plant immune receptors (21, 22).

Chitin, a linear polymer of β-1,4-linked N-acetylglucosaminyl residues, is synthesized by membrane-bound glycosyltransferase (GT) family 2 enzymes called chitin synthases (CHSs) (18, 23). In terms of glucans, the majority is of the β-1,3-linked type, which in some instances is cross-linked with chitin to form the core structure of the fungal cell wall. Additionally, different proportions of branched β-1,6-glucan can be found in some fungi, usually extending to the cell wall surface where it forms connections to mannoproteins (19, 24, 25). β-1,3-glucan is synthesized by a membrane-bound enzyme from GT family 48 (GT48) (25, 26). The enzymes required for β-1,6-glucan biosynthesis have not been described in any filamentous fungal species (27). However, multiple enzymes associated with β-1,6-glucan biosynthesis and β-1,3-glucan modification have been described in the yeast *Saccharomyces cerevisiae* (18, 27, 28).

Given the immunogenic nature of some cell wall carbohydrates, plant-associated fungi must modify their cell wall during host colonization to avoid detection (29, 30). Likewise, as the apoplast is rich in plant-derived glucanases and chitinases (29), fungi must also actively prevent the hydrolytic release of chitin and β-glucan oligomers from their cell walls (31). One proposed strategy used by fungi is to deacetylate chitin to chitosan (32-35), which is a poor elicitor of plant defences (36-38) and a weak substrate of plant chitinases (39, 40). Another strategy is to accumulate α-1,3-glucan on the cell surface, which shields it from the action of plant hydrolases and, in doing so, prevents the release of carbohydrate-based MAMPs (41-44).

In line with the importance of the fungal cell wall, its structure and the enzymes required for its biosynthesis are common targets for antifungal compounds (24, 45). As such, knowledge of fungal cell wall carbohydrate composition is important for the development of novel fungicides and, therefore, control strategies. To date, a detailed analysis of the cell wall carbohydrate composition in a *Venturia* species has not been published, with only one study in 1965 reporting that the cell wall of *V. inaequalis* grown in culture is made up of 28% hemicellulose, 13% β(γ)-cellulose, 20% α-cellulose and 7% chitin (46). As cellulose is generally accepted to be absent from fungal cell walls (47), a more thorough investigation of the cell wall carbohydrate composition in *V. inaequalis* using state-of-the-art techniques is now needed. Here, using a glycosidic linkage analysis, with support from gene expression and proteomic data, we report the cell wall carbohydrate composition of sporulating tubular hyphae from *V. inaequalis* developed on the surface of CMs. Then, using confocal laser scanning microscopy (CLSM), again in conjunction with gene expression data, we provide insights into the cell wall carbohydrate composition of infection structures developed by *V. inaequalis in planta* and compare these to the infection-like structures developed in CMs.

## Results

### The major cell wall polysaccharides of sporulating tubular hyphae formed by *V. inaequalis* in culture are glucans and mannans

To investigate the carbohydrate composition of the *V. inaequalis* cell wall during growth in culture, a glycosidic linkage analysis was performed on cell walls harvested from sporulating tubular hyphae developed on CMs overlaying potato dextrose agar (PDA) (**Figure 1**). The fungal material was extensively washed to avoid contamination from the underlying PDA. The glycosidic linkage analysis revealed that most polysaccharides present in the *V. inaequalis* cell wall were composed of glucosyl (Glc) (~44%) and mannosyl (Man) (~37%) residues, followed by unidentified hexopyranosyl (Hxp) (~10%), galactosyl (Gal) (~8 %) and N-acetylglucosaminosyl (GlcNAc) (~4%) residues (**Figure 2**). This analysis also revealed that the most dominant Glc linkage was 1,3-Glc (41.7%), followed by 1,4-Glc (26%), terminal (t)-Glc (9.63%), 1,3,6-Glc (8.03%), 1,6-Glc (3.8%) and 1,4,6-Glc (3.5%) (**Figure 2**). The entire GlcNAc fraction consisted of 1,4-GlcNAc residues, while the most dominant Man linkage was t-Man (50.4%), followed by 1,2-Man (33.1%). Finally, the Hxp fraction consisted of only two linkages, 2,6-Hxp (78.3%) and 4,6-Hxp (21.7%), while the Gal fraction was mostly t-Gal (87.2%) and 1,4-Gal (12.8%) (**Figure 2**).

**Figure 1.**
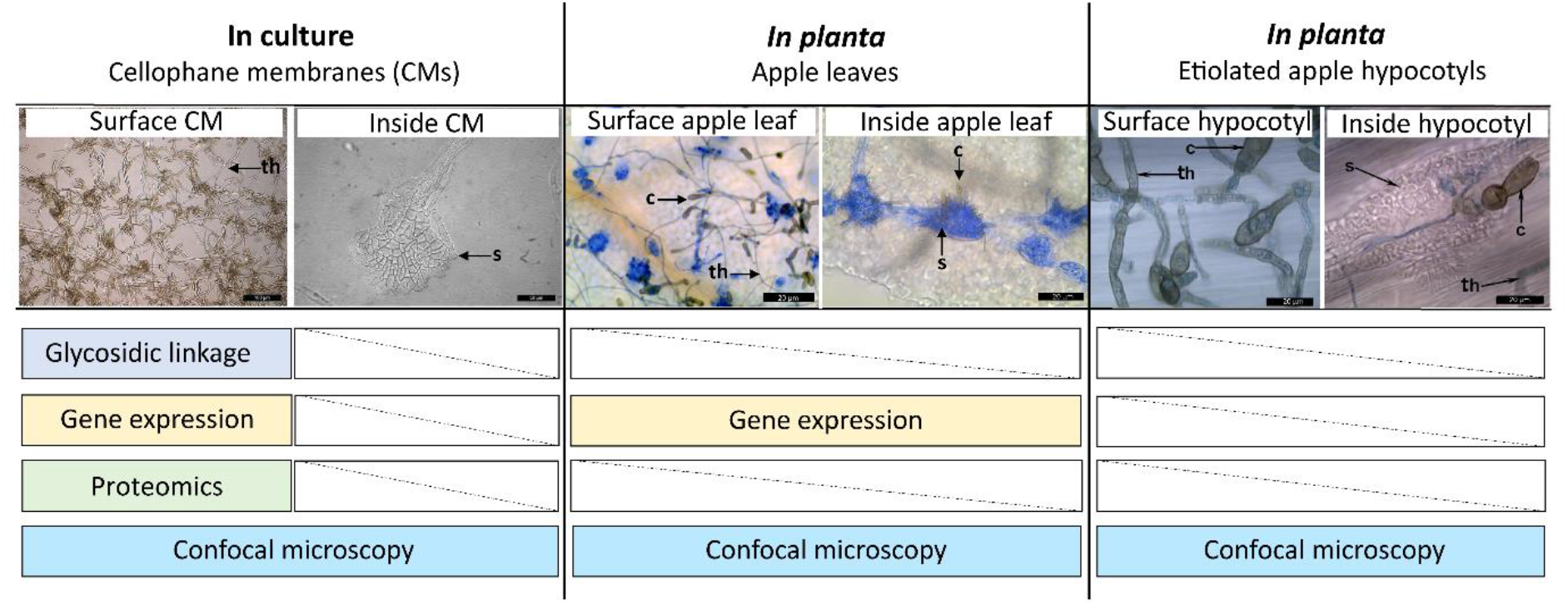
Summary of *Venturia inaequalis* samples used in this study. Tubular hyphae growing on the surface of cellophane membranes (CMs) overlaying potato dextrose agar were used for the glycosidic linkage analysis, proteomic analysis, gene expression (RNA-seq) analysis, as well as confocal laser scanning microscopy (CLSM). Infection-like structures formed inside CMs were used for CLSM. Infected apple leaves were used for the gene expression analysis and CLSM. Infected etiolated apple hypocotyls (a model *in planta* infection system) were used for CLSM. All scale bars: 20 µm. c, conidium; s, stroma; th, tubular hypha.

**Figure 2.**
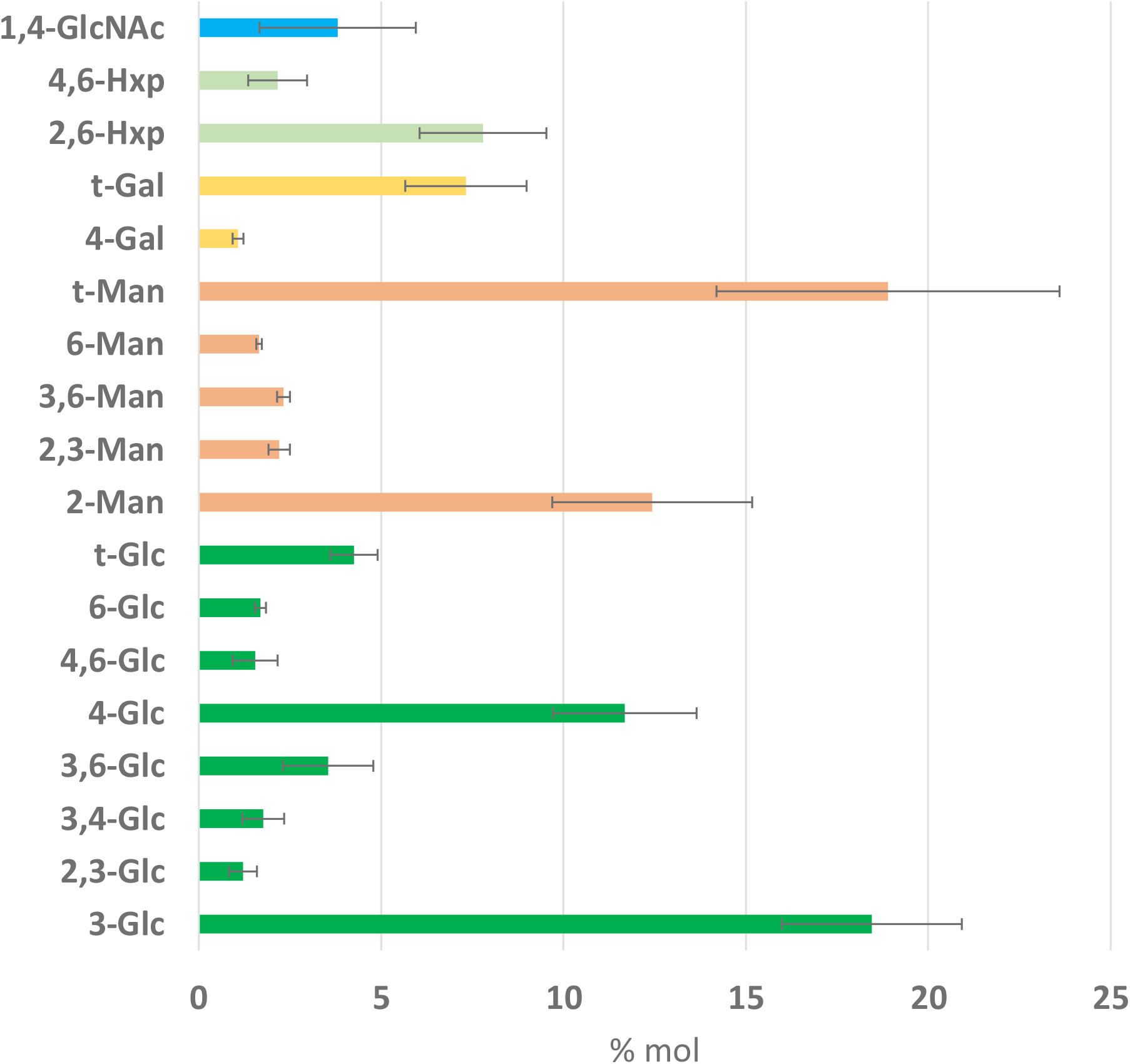
Glycosidic linkage analysis (mole percentage, % mol) of the cell wall carbohydrate fraction from sporulating tubular hyphae of *Venturia inaequalis* developed on the surface of cellophane membranes (CMs) overlaying potato dextrose agar at 5 days post-inoculation. Man, mannose; Glc, glucose; Gal, galactose; Hxp, hexopyranose; GlcNAc, N-acetylglucosamine. Error bars represent standard deviation across three technical replicates.

### Identification and expression of genes putatively associated with cell wall polysaccharide biosynthesis in sporulating tubular hyphae of *V. inaequalis*

To determine which cell wall carbohydrate biosynthetic genes are expressed during growth of *V. inaequalis* on the surface of CMs, we used a combination of bioinformatic, proteomic and transcriptomics approaches (**Figure 1**). As a starting point, the most recently predicted gene catalogue for *V. inaequalis* (48) was inspected to identify genes putatively associated with cell wall biosynthesis (**Table S1, Supplementary file 1**). Based on this analysis, 231 genes were identified (**Supplementary file 1**). Next, the expression of these genes was investigated using pre-existing RNA-seq data from sporulating tubular hyphae of *V. inaequalis* grown on CMs at 7 dpi (48). This analysis revealed that, of the 231 predicted cell wall biogenesis genes, 135 were expressed with a transcripts per million (TPM) value >10 (**Figure 3**). Finally, a proteomic analysis of total protein from sporulating tubular hyphae of *V. inaequalis* grown on CMs at 5 dpi was performed. Based on this analysis, 24 of the 231 putative cell wall biosynthetic genes were found to encode proteins with proteomic support (**Supplementary file 3** and **4**), confirming that they were indeed produced. Of these, 17 had a TPM expression value >10 (**Figure 3**).

**Figure 3.**
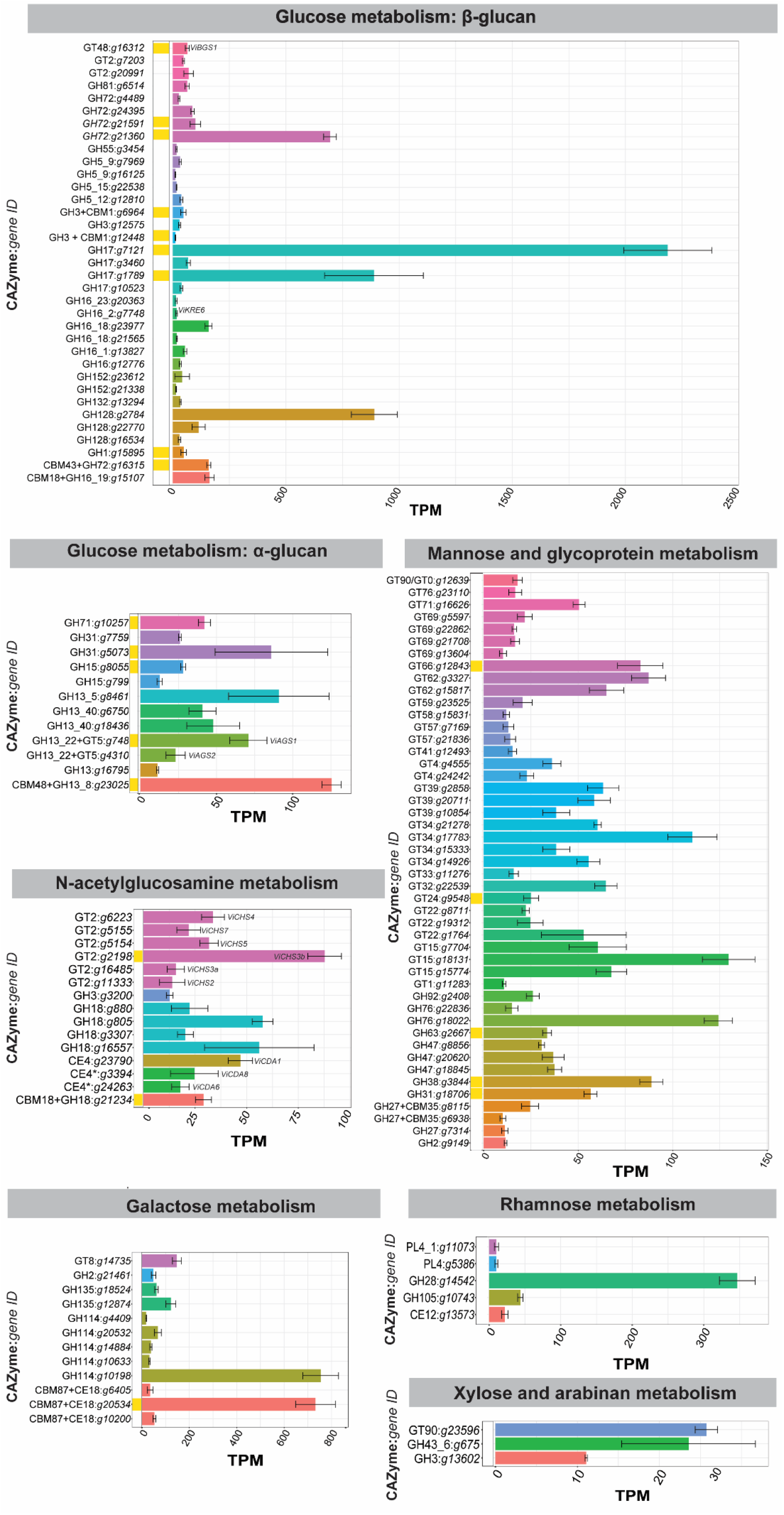
Expression of carbohydrate-active enzyme (CAZyme)-encoding genes from *Venturia inaequalis* putatively associated with cell wall biosynthesis during growth as sporulating tubular hyphae on the surface of cellophane membranes overlying potato dextrose agar at 7 days post-inoculation. Gene expression data are transcripts per million (TPM), averaged from four biological replicates, with error bars representing standard deviation. Only genes with TPM values >10 are shown and are grouped into families by colour. Enzymes labelled with an asterisk (*) were identified by Protein family (Pfam) search. Yellow blocks indicate proteins that have proteomic support by mass spectrometry. AGS, α-1,3-glucan synthase; BGS, β-1,3-glucan synthase; CBM, carbohydrate-binding module; CDA, chitin deacetylase; CE, carbohydrate esterase; CHS, chitin synthase; GH, glycoside hydrolase; GT, glycosyltransferase; PL, polysaccharide lyase.

Among the glycosyltransferases (GTs) identified, 13 were putative family 2 enzymes and, of these, eight were annotated as CHSs. These were named ViCHS1–7 according to the previously established CHS classification scheme (49), based on both their CHS domain (**Figure 4.A**) and phylogenetic distribution (**Figure 4.B**). *V. inaequalis* had at least one representative from each CHS class (class I–VI), and two CHSs from class III (**Figure 4.A**). Four CHSs were from division I, having a simple ‘amino (N)-terminus CHS domain 1’ plus ‘CHS domain 1’ structure (**Figure 4.A**). All division I CHSs, except ViCHS1, were encoded by genes that had a TPM expression value >10, with *ViCHS3b* being the most highly expressed and the only CHS with proteomic support (**Figure 3**). Three CHSs were from division II and, of these, ViCHS4 only had a single ‘CHS domain 2’ module (**Figure 4.A**). In contrast, ViCHS5 and ViCHS7 both had a ‘cytochrome b5-binding domain’ and a ‘Dek domain’ at their carboxyl (C)-terminus, while ViCHS5 also had an N-terminal ‘myosin motor-like’ domain (**Figure 4.A**). Finally, ViCHS6 was from division III and had a single C-terminal ‘CHS domain 2’ module (**Figure 4.A**). The gene encoding this CHS, however, was not highly expressed in culture (<10 TPM). All CHSs contained both a QxxRW motif required for catalytic activity, as well as QxxEY and EDRxL domains of unknown function (**Figure S1**) (23).

**Figure 4.**
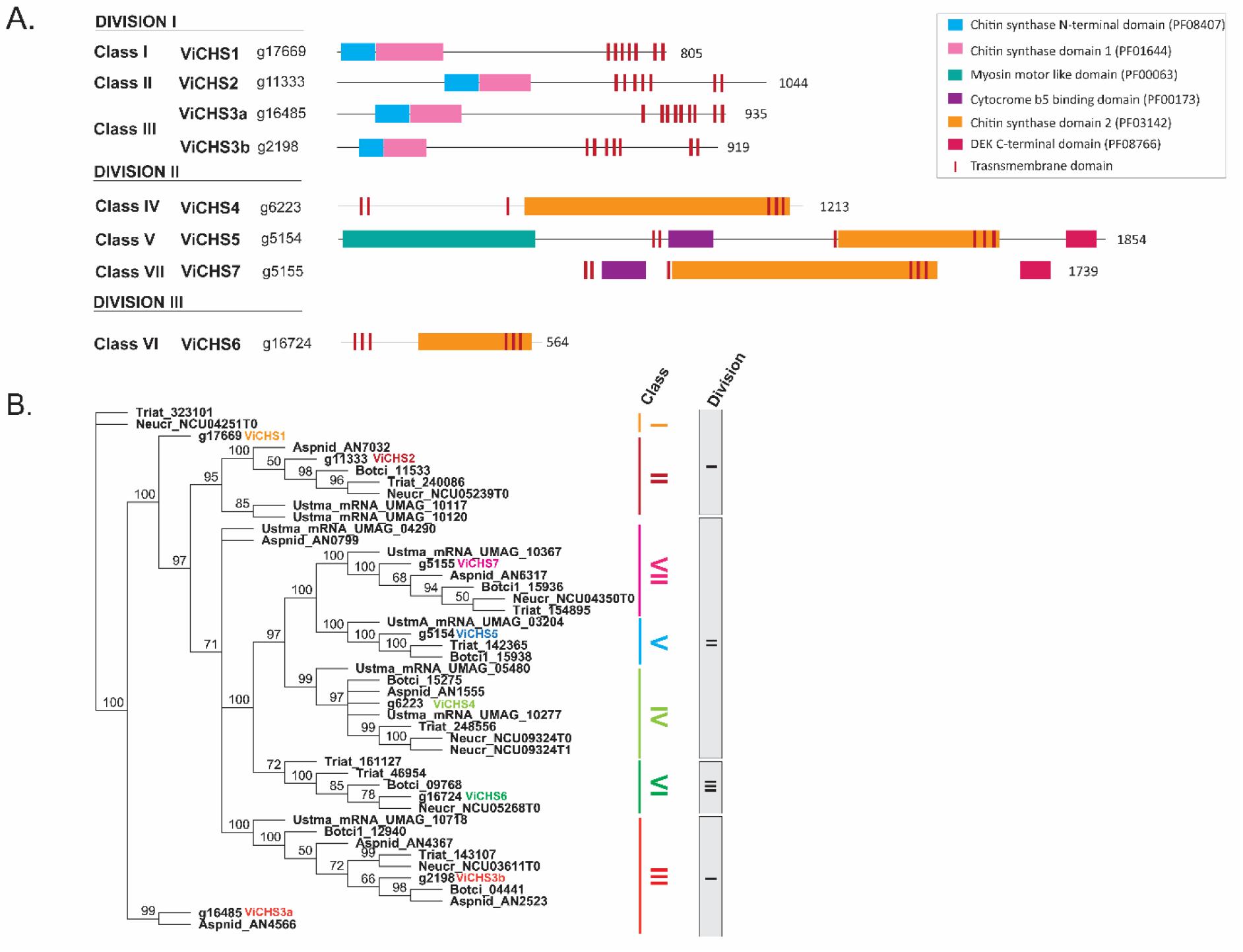
Chitin synthase (CHS) proteins of *Venturia inaequalis*. **A**. Predicted classification and domain organization of CHS proteins. **B**. Phylogenetic classification of the eight predicted CHSs. CHSs from *Aspergillus nidulans* (Aspnid), *Neurospora crassa* (Neucr), *Botrytis cinerea* (Botci) and *Ustilago maydis* (Ustma) were included for reference. *V. inaequalis* CHS proteins are highlighted in bold and coloured letters. Phylogenetic tree was generated from a MUSCLE alignment and end-joining method using Geneious v9.0.5, with node values indicating consensus support (%) from 100 replicates.

Multiple enzymes putatively associated with chitin modification were encoded by genes expressed during growth of *V. inaequalis* in culture, including five glycoside hydrolase (GH) family 18 chitinases, and one β-N-acetyl hexosaminidase of GH family 3 (**Figure 3, Supplementary file 1**). However, only one chitinase had proteomic support (**Figure 3**). In total, eight putative chitin deacetylase (*CDA*) genes were identified (**Supplementary file 1**). Of those (**Figure S2.A**), a sequence alignment revealed that only ViCDA1 and ViCDA7 possessed all previously described conserved residues for catalytic activity and metal-binding and were likely functional (50, 51) (**Figure S2.B**). ViCDA4 had all metal-binding and catalytic site residues, except one, as the second conserved aspartic acid for catalytic activity was substituted by a similar negatively charged amino acid, glutamic acid. Therefore, it is likely that ViCDA4 is functional. ViCDA5 and ViCDA6 had all catalytic site residues, except the last histidine, and an amino substitution in the first metal-binding position (**Figure S2.B**). As such, these two proteins might not be functional. ViCDA3 had two amino acid substitutions at the first and third metal-binding positions which are characteristic of allantoinases; enzymes that hydrolyse allantoin, a nitrogen-rich organic compound (52) (**Figure S2.B**). Therefore, ViCDA3 might function as an allantoinase instead of a CDA. Finally, due to a C-terminal truncation, ViCDA8 was missing the last two catalytic site residues and, therefore, is likely to be non-functional (**Figure S2.B**). None of the CDAs had a predicted transmembrane domain or glycosylphosphatidylinositol (GPI) anchor, while only ViCDA1 and ViCDA4 had a predicted N-terminal signal peptide for secretion (**Figure S2.A**). *ViCDA1* was the only potentially active CDA-encoding gene with a high level of expression during growth of *V. inaequalis* in culture; however, this enzyme did not have proteomic support (**Figure 3 and Figure S2.C**).

Only one gene encoding a putative β-1,3-glucan synthase of GT family 48, *ViBGS1*, was identified in the *V. inaequalis* genome (**Supplementary file 1**). This gene was expressed in culture and the resulting enzyme was detected by the proteomic analysis (**Figure 3**). Additionally, many genes encoding β-1,3-glucan-modifying enzymes were identified, such as members of the GH17 and GH72 families (**Supplementary file 1**), which are anticipated to assist in either the elongation or branching of β-1,3-glucans (28, 53). Of these, four *GH17* and four *GH72* genes were expressed during growth of *V. inaequalis* in culture and half of these had proteomic support (**Figure 3**). This included the *GH17* gene *g7121*, which was the most highly expressed carbohydrate-active enzyme (CAZyme)-encoding gene.

A further 13 genes encoding putative GH family 16 enzymes were also identified (**Supplementary file 1**), with four of these found to be expressed during growth in culture (**Figure 3**). Three of these possibly encode chitin transglycosylases required to cross-link chitin with glucan (54), while the fourth encodes a KRE6-like enzyme (ViKRE6) that is possibly associated with β-1,6-glucan biosynthesis (55, 56). None of the four GH16 enzymes were identified in the proteomic analysis.

The genome of *V. inaequalis* also carried two genes encoding putative α-1,3-glucan synthases, named *ViAGS1* and *ViAGS2* (**Supplementary file 1**). Both were expressed in culture; however, only ViAGS1 had proteomic support (**Figure 3**). Finally, the *V. inaequalis* genome possessed 71 genes involved in the biosynthesis of mannan (**Supplementary file 1**). Of these, six encoded mannan polymerases and 25 encoded mannosyltransferases expressed with a TPM value >10 in culture. However, none of these enzymes had proteomic support (**Figure 3**).

### Genes putatively associated with the biogenesis of carbohydrate-based MAMPs from *V. inaequalis* are down-regulated during host colonization

A glycosidic linkage analysis could not be performed to determine the carbohydrate composition of infection structures produced by *V. inaequalis* during infection of apple tissue. This was due to the paucity of fungal material generated during subcuticular growth. Thus, to make inferences about how the *V. inaequalis* cell wall carbohydrate composition changes during host infection, relative to growth in culture, the expression of the 231 putative cell wall biosynthesis genes identified above was investigated using pre-existing *in planta* transcriptomic data (48) (**Figure 1**). These data were collected from apple leaves infected with *V. inaequalis* at six time points: 12 and 24 hours post-inoculation, as well as 2, 3, 5 and 7 dpi (48). Using these data, a total of 68 genes putatively associated with fungal cell wall biosynthesis were found to be up-regulated, and 43 down-regulated, at one or more *in planta* time points compared to growth in culture (**Supplementary file 2**). Interestingly, most differentially expressed genes were associated with β-glucan metabolism (**Figure 5.A** and **Figure S3**). More specifically, during early infection (12–24 hpi), the *Gas5/Gel1-like* gene, which encodes a GH72 1,3-β-glucanosyltransferase with sequence similarity to Gas5 enzymes from yeast and Gel1 from *Aspergillus* spp. (57, 58), was up-regulated. Later, during mid-late infection (5 and 7 dpi), several β-glucosidase-encoding genes were up-regulated. Only a few genes associated with β-glucan metabolism were down-regulated during host colonization, such as two genes encoding GH17 proteins and the KRE6-like enzyme (**Figure 5.A** and **Figure S3**). Regarding α-1,3-glucan metabolism, *ViAGS2* was up-regulated during early (12 and 24 hpi) host infection (**Figure 5.B**).

**Figure 5.**
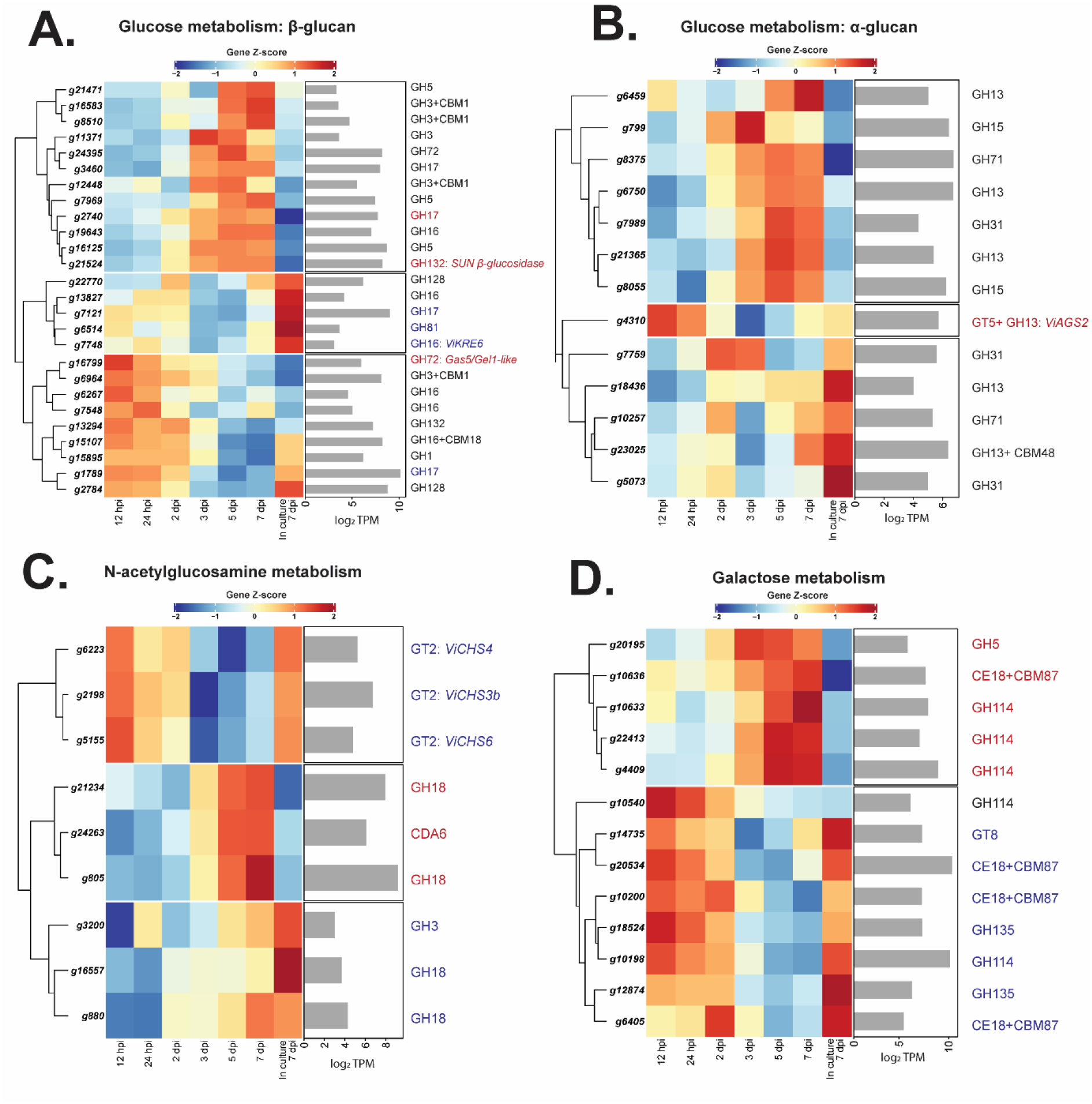
Heatmaps showing the expression profiles of genes from *Venturia inaequalis* that are both putatively associated with cell wall biosynthesis and differentially expressed during host colonization, relative to growth in culture. Only differentially expressed genes putatively associated with **A**. glucose (β-glucan), **B**. glucose (α-glucan), **C**. N-acetylglucosamine, and **D**. galactose metabolism at one or more *in planta* time points when compared to growth on cellophane membranes overlying potato dextrose agar, and with a minimum expression of 10 transcripts per million (TPM) are shown. Heatmaps are rlog-normalized counts across all samples (Z-score), averaged from four biological replicates. Bar plots depict the maximum log_2_ TPM count value across all *in planta* time points. hpi, hours post-inoculation; dpi, days post-inoculation; CBM, carbohydrate-binding module; CDA, chitin deacetylase; CE, carbohydrate esterase; CHS, chitin synthase; GH, glycoside hydrolase; GT, glycosyltransferase. Blue colour highlight genes down-regulated and red colour highlights genes up-regulated during host-colonization, relative to growth in culture.

Genes involved in chitin metabolism were, in general, down-regulated during host colonization (**Figure 5.C** and **Figure S4**). During early infection, two genes encoding putative GH18 chitinases and one gene encoding a GH3 enzyme of unknown function, were down-regulated. Later, during mid-late infection, the *CHS* genes *ViCHS3b, ViCHS4* and *ViCHS7* were down-regulated. Interestingly, from 3 dpi, two chitinase-encoding genes, as well as the gene encoding the putatively inactive ViCDA6 were up-regulated (**Figure 5.C** and **Figure S4**). Additionally, *ViCDA1*, which encodes the putatively secreted and active CDA, was constitutively expressed both in culture and *in planta* (**Figure S2.C**).

Several genes associated with galactose metabolism were down-regulated during mid-late infection. However, others were up-regulated over this infection stage, especially three genes encoding GH114 proteins (**Figure 5.D**). Finally, most genes associated with mannose and glycoprotein metabolism were up-regulated, especially those genes encoding putative mannosidases (**Supplementary file 2**).

### Chitosan is present on the surface of *V. inaequalis* infection structures formed *in planta*, while chitin is restricted to septa

To further investigate the cell wall carbohydrate composition of the cellular morphotypes formed by *V. inaequalis*, we used CLSM in conjunction with different carbohydrate-specific probes and antibodies to label chitin, chitosan, β-1,3-glucan, and α-1,3-glucan. CLSM was performed on *V. inaequalis* growing in culture on the surface and inside CMs, as well as during host colonization on the surface and inside living host tissue (**Figure 1**). For host colonization, two approaches were taken, with the first involving detached apple leaves, and the second involving detached etiolated apple hypocotyls (**Figure 1**). Here, etiolated apple hypocotyls were included as they have previously been shown to be a good system for visualising infection by *V. inaequalis* due to their reduced tissue thickness, as well as their lower levels of chlorophyll, other pigments and phenolic compounds, when compared to apple leaves (59). However, it is important to note that hypocotyl infection does not occur in orchards, as these apples are cultivated from clonal bud wood, not seed. Hypocotyl infection may, though, occur in natural apple forests.

To monitor the distribution of chitin on the surface of the fungal cell wall, all samples were labelled with wheat germ agglutinin (WGA) conjugated to the fluorophore AlexaFluor (AF) 488 (WGA^488^) (**Figure 6** and **Figure S6**). Tubular hyphae formed on the surface of CMs (**Figure 6.A**), infected apple leaves (**Figure 6.D,E**) and etiolated hypocotyls (**Figure S6**) presented limited amounts of surface-exposed chitin, with labelling mainly restricted to hyphal breakage points and, sometimes, septa. The maximum fluorescence intensity occurred at the cell periphery, indicating that the labelling signal was derived from the cell wall (**Figure 6.B**). To label the total chitin present in the cell wall, fungal material was first permeabilized with NaOH and then labelled with the chitin stain calcofluor white. By doing so, faint labelling was observed around the periphery of tubular hyphae, with stronger labelling found at septa (**Figure 6.C**). On the infection-like structures developed inside CMs, chitin labelling was observed around the periphery of runner hyphae, as well as at the septa of runner hyphae and stromata (**Figure 6.A**). In contrast, following host penetration, chitin was exclusively restricted to septa (**Figure 6** and **Figure S6**).

**Figure 6.**
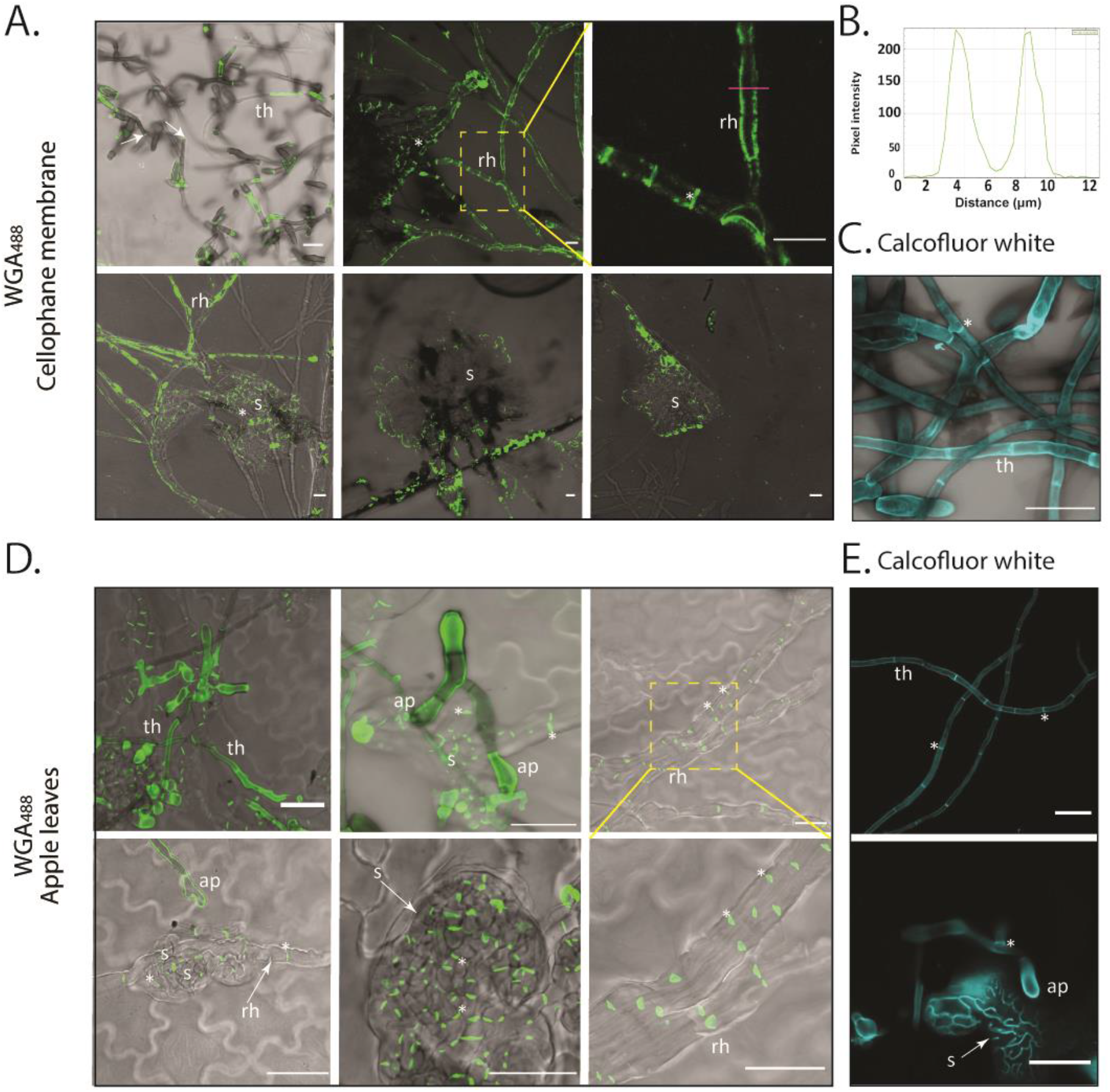
Labelling of chitin on *Venturia inaequalis* cellular morphotypes developed in culture and *in planta*. **A**. Cellular morphotypes developed in culture in and on cellophane membranes (CMs) overlaying potato dextrose agar labelled with wheat germ agglutinin-AlexaFluor 488 (WGA^488^) (green pseudocolour). **B**. Plot profile of pixel intensity along a line (magenta) in A (top right-hand image) made with ImageJ 1.x. **C**. Tubular hyphae developed on a CM labelled with calcofluor white (cyan pseudocolour) after permeabilization with NaOH. **D**. Cellular morphotypes developed in and on ‘Royal Gala’ apple leaves labelled with WGA^488^ (green pseudocolour). **E**. Tubular hyphae developed on ‘Royal Gala’ apple leaves labelled with calcofluor white (cyan pseudocolour) after permeabilization with NaOH. Dashed yellow squares indicate zoomed-in areas. ap, appressorium; rh, runner hyphae; s, stroma; th, tubular hypha; arrow, hyphal breakage; *, septum. All scale bars: 20 μm.

Next, as *V. inaequalis* has eight putative *CDA* genes (**Figure 5**) and, of these, *ViCDA6* is up-regulated during infection, while *ViCDA1* is constitutively expressed in culture and *in planta*, it was hypothesized that *V. inaequalis* deacetylates chitin to chitosan during host colonization. Therefore, the tubular hyphae and infection structures of *V. inaequalis* developed in culture and *in planta* were probed using the oligogalacturonate (OGA) probe conjugated with AF488 (OGA^488^) (60).

Chitosan labelling was not detected on tubular hyphae developed on the surface of CMs or apple leaves (**Figure 7.A, C**). In contrast, tubular hyphae developed on etiolated hypocotyls had surface-exposed chitosan (**Figure 7.D**), highlighting a key difference among these host infection systems. The reason behind chitosan induction on the surface of etiolated hypocotyls is not clear, but we hypothesize that *V. inaequalis* may grow more intimately with the cutin layer and, as part of this, there might be some plant-derived trigger (e.g. cutin monomers) inducing chitosan production. With this in mind, we tested if chitosan production could be induced in culture. For this purpose, wax was extracted from apple fruit and added to the surface of CMs before inoculation with *V. inaequalis* conidia. Remarkably, following germination, apple wax triggered chitosan production on tubular hyphae developed in culture (**Figure 7.E**). The maximum fluorescence intensity from chitosan labelling occurred at the cell periphery, indicating that the labelling signal was derived from the cell wall (**Figure 7.B**). Regarding the infection structures developed *in planta*, as well as the infection-like structures developed in CMs, chitosan labelling was also observed at the periphery (**Figure 7.C, D**).

**Figure 7.**
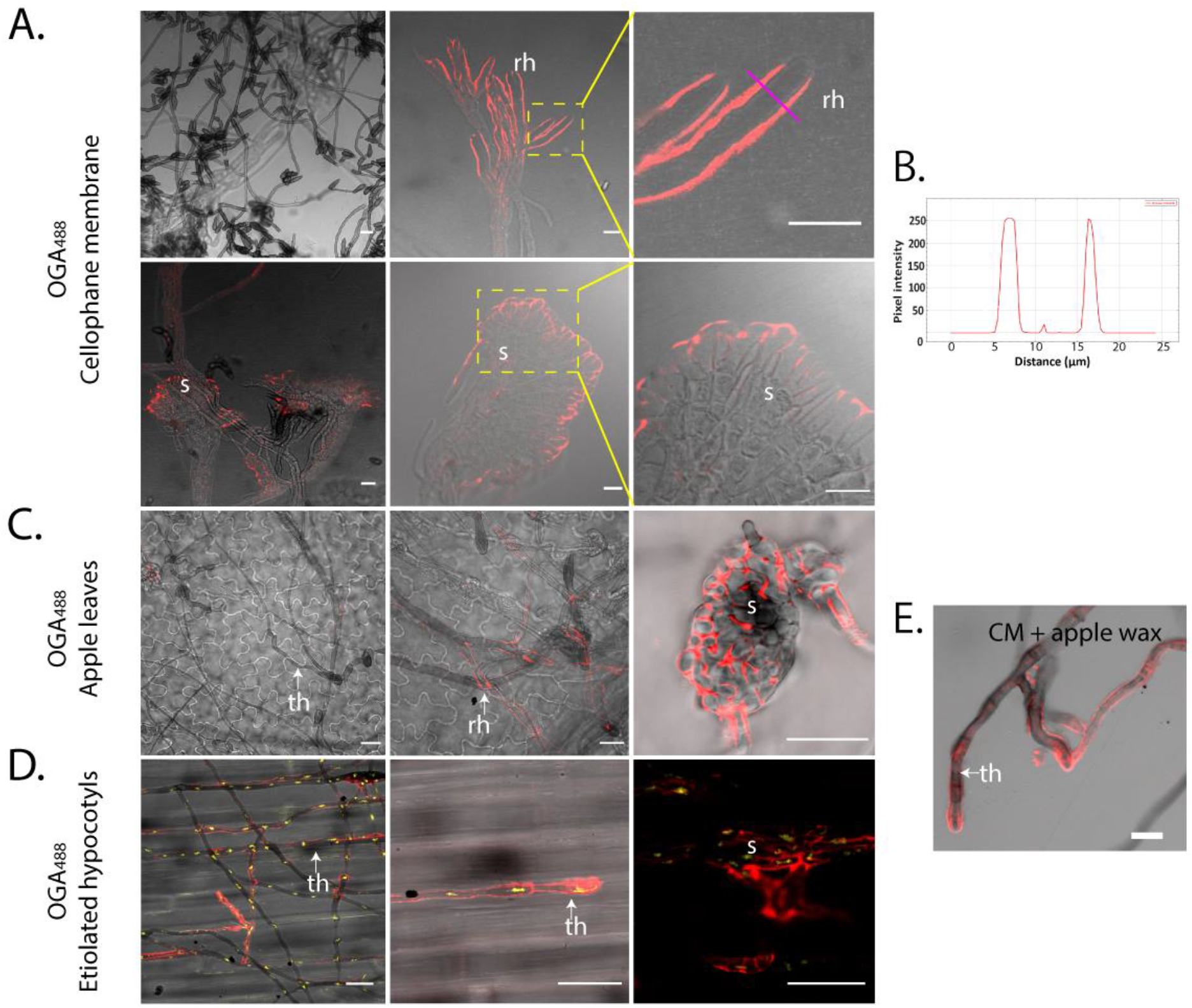
Label-accessible chitosan on the surface of *Venturia inaequalis* cellular morphotypes developed in culture and *in planta*. The fluorophore-labelled oligosaccharide OGA^488^ was used to visualize chitosan (red pseudocolour). **A**. *V. inaequalis* cellular morphotypes developed in culture in and on cellophane membranes (CMs) overlying potato dextrose agar. **B**. Plot profile of pixel intensity along a line (magenta) in A (top right-hand image) made with ImageJ 1.x. **C**. Cellular morphotypes developed in and on apple leaves. **D**. Cellular morphotypes developed in and on etiolated hypocotyls, fungal nuclei labelled with propidium iodide (PI, yellow pseudocolour). **E**. Surface hyphae developed on a CM covered with apple wax. All scale bars: 20 μm. ap, appressorium; rh, runner hyphae; th, tubular hypha; s, stroma. Dashed yellow squares indicate zoomed-in areas.

Next, we investigated the distribution of β-1,3-glucan during growth of *V. inaequalis* in culture and *in planta*, as β-1,3-glucan is known to be an elicitor of plant defences (61). For this purpose, β-1,3-glucan localization was investigated with a primary mouse antibody specific to β-1,3-glucan in conjunction with the anti-mouse secondary antibody CF-488. β-1,3-glucan-specific labelling was rarely observed around tubular hyphae developed on the surface of CMs, with labelling mostly observed at hyphal breakage points (**Figure 8.A**). To label β-1,3-glucan present on the surface of infection-like structures formed inside CMs, sandpaper had to be used to create antibody entry points. In doing so, we observed an intense but patchy labelling at the periphery of the infection-like structures (**Figure 8.A**). A similar patchy distribution was observed on tubular hyphae developed on the surface of apple leaves and etiolated hypocotyls (**Figure 8.B and Figure S6.B**). Interestingly, labelling of β-1,3-glucan was not observed on young infection structures developed in leaves (**Figure 8.B**) or hypocotyls (**Figure S6.B**). Instead, labelling of β-1,3-glucan could only be observed on mature stromata that had ruptured the apple cuticle upon sporulation (**Figure 8.B**).

**Figure 8.**
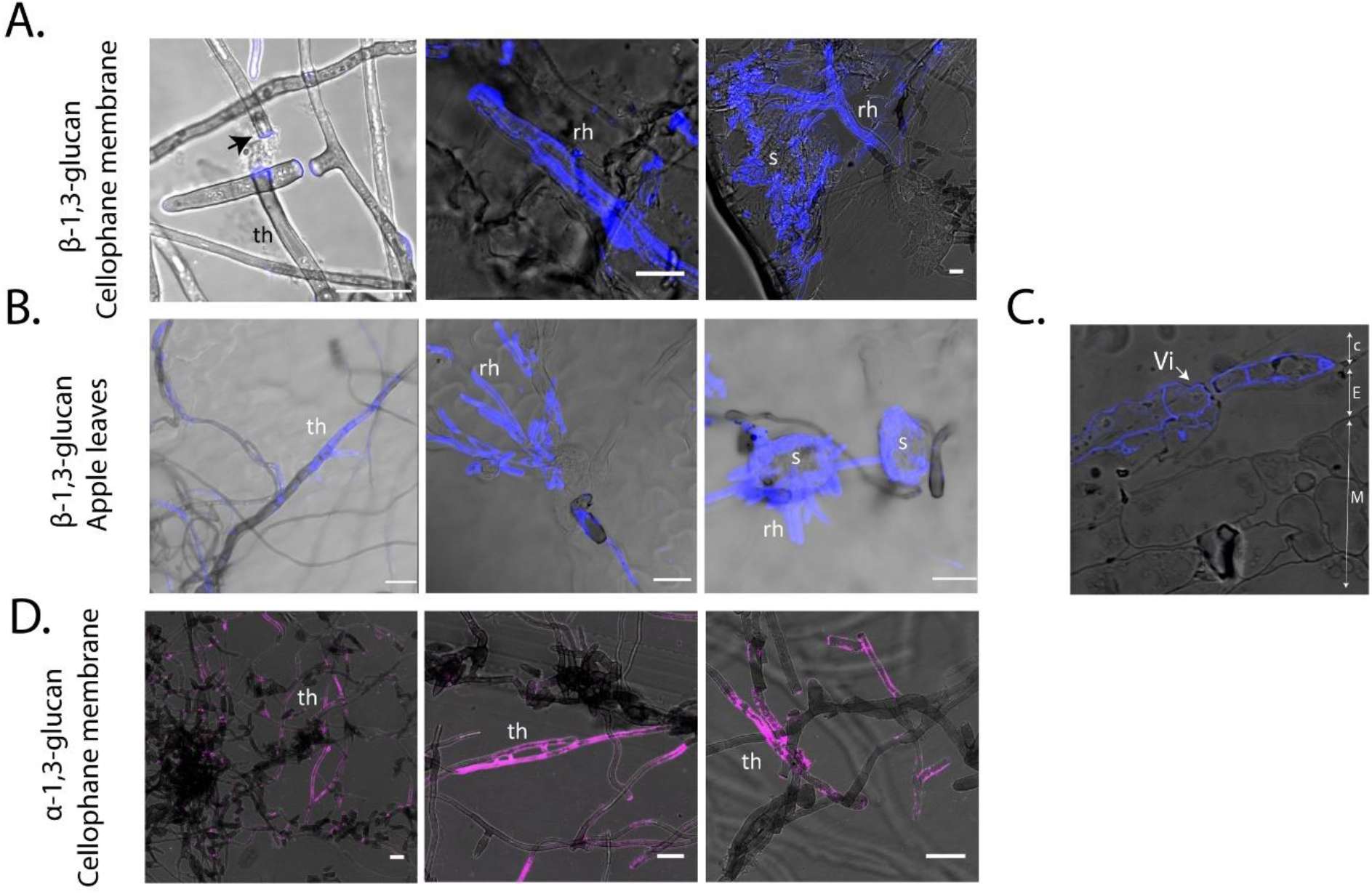
Label-accessible β-1,3-glucan and α-1,3-glucan of *Venturia inaequalis* (Vi) cellular morphotypes developed in culture and *in planta*. The monoclonal anti β-1,3-glucan primary antibody and CF-488 secondary antibody were used to label β-1,3-glucan (blue pseudocolour), while the monoclonal MOPC-104E primary antibody and CF-488 secondary antibody were used to label α-1,3-glucan (pink pseudocolour). **A**. β-1,3-glucan labelling of *V. inaequalis* cellular morphotypes developed in culture in and on cellophane membranes (CMs). **B**. β-1,3-glucan labelling of *V. inaequalis* cellular morphotypes developed in and on detached apple leaves. **C**. β-1,3-glucan labelling of a cross-section of a detached leaf infected with *V. inaequalis*. C: apple cuticle, E: apple epidermal cells, M: apple mesophyll cells. **D**. α-1,3-glucan labelling of *V. inaequalis* cellular morphotypes developed in culture in and on CMs. All scale bars: 20 μm. Rh, runner hyphae; s, stroma; th, tubular hyphae; arrow: hyphal breakage points.

As *V. inaequalis* has two putative α-1,3-glucan synthase genes and, of these, *ViAGS2* is up-regulated during early host colonization (**Figure 5**), we attempted to label α-1,3-glucan using the primary antibody MOPC-104E in conjunction with the secondary antibody anti-mouse CF-488. Labelling of α-1,3-glucan was observed on tubular hyphae developed on the surface of CMs (**Figure 8.D**), but not on tubular hyphae developed on the surface of apple tissue (data not shown). Likewise, no labelling of α-1,3-glucan was observed on the surface of infection-like structures formed in culture or on the infection structures developed *in planta* (data not shown).

## Discussion

Even though the cell wall plays a crucial role in viability and pathogenesis (20, 30, 62), few studies have focused on the cell wall of plant-pathogenic fungi. Here, we examined the cell wall carbohydrate composition of sporulating tubular hyphae from *V. inaequalis* developed on the surface of CMs using glycosidic linkage analysis. As observed in other fungi, the most abundant glucosyl linkages found in *V. inaequalis* were 1,3-Glc, with the proportion identified similar to that found in the subcuticular plant pathogen, *R. secalis* (**Table 1**) (63). Glycosidic linkage analysis does not allow discrimination between α- and β-linkages. However, using CLSM, we were able to show that α-1,3-glucan was present on surface hyphae developed in culture. Therefore, a fraction of the identified 1,3-Glc could be from α-1,3-glucan, which coats the surface of the *V. inaequalis* cell wall.

**Table 1.** Carbohydrate composition (%) of the *Venturia inaequalis* cell wall, determined in this study using glycosidic linkage analysis, compared to other fungal species for which carbohydrate composition is known. This comparison was made for the filamentous fungi *Blumeria graminis* f. sp. *hordei* (67), *Rhynchosporium secalis* (63), *Neurospora crassa* (68), *Aspergillus fumigatus* (69) (alkali-insoluble fraction of the cell wall), and the yeast *Saccharomyces cerevisiae* (70). This table was extracted and modified from (67). Gal, galactose; Glc, glucose; GlcNAc, N-acetylglucosamine; Man, mannose; Hxp, hexopyranose.

|  | *V. inaequalis* | *B. graminis*<br>f.sp. *hordei* | *R. secalis* | *N. crassa* | *A. fumigatus* | *S. cerevisiae* |
| --- | --- | --- | --- | --- | --- | --- |
| t-Man | 18.9 | 1.7 | 3.1 | 1.5 | 0.1 | 15.7 |
| t-Glc | 4.3 | 5.7 | 13.9 | 13 | 0.3 | 5.5 |
| t-Gal | 7.3 | 2.2 | 0 | 1 | 2.2 | 0 |
| 3-Glc | 18.5 | 35.6 | 11.6 | 54 | 43.3 | 26.9 |
| 2-Man | 12.4 | 4.3 | 4.3 | 3 | 2.4 | 7.9 |
| 6-Man | 1.7 | 3.8 | 4.4 | 0 | 1.2 | 1.2 |
| 6-Glc | 1.7 | 2 | 0.5 | 0 | 0.3 | 0.7 |
| 4-Gal | 1.1 | 6.8 | 0 | 0 | 0 | 0 |
| 4-Glc | 11.7 | 6.3 | 17.2 | 4 | 14.2 | 0 |
| 2,3-Man | 2.2 | 0 | 0.9 | 0 | 0 | 0 |
| 3,4-Glc | 1.8 | 2.1 | 0.5 | 0 | 0.3 | 0.7 |
| 2,3-Glc | 1.2 | 7.5 | 1.5 | 0 | 0 | 7 |
| 3,6-Glc | 3.5 | 1.6 | 2.7 | 5 | 1.6 | 2 |
| 3,6-Man | 2.3 | 0.6 | 0.6 | 0 | 0 | 0.4 |
| 2,6-Hxp | 7.8 | 0 | 0 | 0 | 0 | 0 |
| 4,6-Glc | 1.5 | 2.3 | 0.5 | 0 | 0 | 0 |
| 4,6-Hxp | 2.2 | 0 | 0 | 0 | 0 | 0 |
| 1,4-GlcNAc | 3.8 | 8.7 | 6.3 | 10 | 17.65 | 0.6 |
| 3,6-Gal | 0.0 | 0 | 0 | 0 | 0 | 0 |

Surprisingly, α-1,3-glucan could not be detected on the surface of *V. inaequalis* infection-like structures developed inside CMs or on the infection structures developed *in planta*, even though the α-1,3-glucan synthase gene, *ViAGS2*, was up-regulated during early host colonization. The lack of α-1,3-glucan labelling *in planta* could indicate that this carbohydrate is not label-accessible on the cell wall surface during host colonization. However, it is more plausible that this is due to the problem we encountered with antibody penetration inside host tissue. Interestingly, in some human fungal pathogens, such as *Histoplasma capsulatum* and *Aspergillus fumigatus*, as well as the fungal plant pathogen *Magnaporthe oryzae*, α-1,3-glucan accumulates on the surface of the fungal cell wall to conceal cell wall-derived MAMPs and, thus, prevents the induction of host defence responses during host infection (41-44, 64). Additionally, in *A. fumigatus*, α-1,3-glucan is important for hyphal and conidial aggregation (65, 66). With this research in mind, additional experiments are now needed to investigate whether α-1,3-glucan is present on the surface of *V. inaequalis* infection structures.

Another abundant glucosyl linkage identified as part of our glycosidic linkage analysis was 1,4-Glc. It cannot be ruled out that a portion of this 1,4-Glc arose from cellulose contamination from the CM itself, or an intracellular form of a glycogen/starch-like polysaccharide that co-purifies with the cell wall during sample preparation. Nevertheless, the low amount of 1,4,6-Glc identified suggests that not all of the 1,4-Glc originated from glycogen/starch-like polymers. Notably, there is very little evidence for the presence of β-1,4-linked glucosyl residues in fungal cell walls although, interestingly, these residues were recently reported in the cell wall of *A. fumigatus* by solid-state nuclear magnetic resonance (47, 71). Hence, it cannot be ruled out that a small fraction of the reported 1,4-Glc from *V. inaequalis* originates from β-1,4-glucan. Another more likely possibility is that a portion of the identified 1,4-Glc forms part of nigeran, a polymer commonly found in Ascomycota fungal cell walls that are composed of α-1,3- and α-1,4-glucans (72).

The presence of 1,3,6-Glc suggests that a fraction of the identified 1,3-Glc is branched. A set of *GH17* and *GH72* genes found to be highly expressed in culture likely encode the enzymes responsible for the presence of 1,3,6-linked glucosyl residues in the *V. inaequalis* cell wall, as these enzymes are putative β-1,3-glucan transglycosylases involved in cross-linking β-1,3-glucans through β-1,6-linkages (28, 53, 54, 73). Interestingly, during mid-late infection, *V. inaequalis* was observed to down-regulate a gene encoding a KRE6 enzyme, *ViKRE6*, which is putatively associated with β-1,6-glucan biosynthesis (55, 56, 74). We suggest that, as reported in *Colletotrichum graminicola* (74), *V. inaequalis* down-regulates *ViKRE6* to reduce surface-exposed β-1,3-6-glucans that would otherwise elicit plant immune responses.

The glycosidic linkage analysis also revealed a relatively large proportion of Man (~37%) and Gal (~8%) residues in the cell wall of *V. inaequalis*, with the majority being terminal. Due to this terminal nature, we cannot conclude from which polymer these residues are derived. The large amount of t-Man (18.9%), however, is similar to the amount of t-Man reported in *S. cerevisiae* (15.7%) (70) and drastically more than other filamentous fungi (1–3%) (63, 67, 68).

Following glycosidic linkage analysis, we attempted to investigate the presence and location of β-1,3- and branched β-1,3-6-glucans on the fungal cell wall surface using a specific antibody that labels β-1,3-glucan. Regarding the infection structures developed inside the host, we only observed strong labelling on the surface of mature stromata that were rupturing through the apple cuticle as part of the sporulation process. It is unclear whether these infection structures were labelled because the mature stromata were more accessible to the β-1,3-glucan antibody, or because the β-1,3-glucan is masked during early host colonization and is only surface-exposed upon sporulation.

The glycosidic linkage analysis also revealed that the cell wall of sporulating tubular hyphae from *V. inaequalis* is comprised of 3.8% chitin, which is relatively low for a filamentous fungus. Indeed, chitin usually makes up around 10–15% of the fungal cell wall (18, 25, 27) (**Table 1**). This result is in line with the labelling profile of chitin present on the surface of tubular hyphae formed on CMs using the WGA^488^ probe and calcofluor white, where chitin was mostly restricted to septa. Transcriptomic and proteomic data obtained in our study suggest that ViCHS3b, a class III CHS enzyme, might be responsible for the bulk of chitin biosynthesis in *V. inaequalis* tubular hyphae.

Interestingly, on the plant surface, chitin was exclusively observed on tubular hyphae and appressoria, while on the infection structures produced after host penetration, chitin was found to be restricted to the septa. Furthermore, genes encoding CHSs and some chitinases were down-regulated during early host colonization. These findings suggest that, in addition to transcriptionally regulating chitin production, *V. inaequalis* also restricts the amount of chitin it exposes on its surface to prevent activation of chitin-triggered host defences. Another possibility is that *V. inaequalis* masks chitin on the cell wall surface through the secretion of chitin-binding effectors, similar to that shown for the Avr4 effector protein of *Fulvia fulva* (75). Alternatively, *V. inaequalis* may produce chitin-binding effectors that sequester chitin oligomers during the process of cell wall remodelling to prevent their detection by the apple immune system. Notably, during *in planta* host colonization, the lysin motif (LysM) domain-containing effector candidate, ViEcp6, which has sequence similarity to the chitin-binding Ecp6 effector of *F. fulva* (76, 77), was found to be up-regulated (48) and had proteomic support in culture (**Supplementary file 3** and **4**). Although Ecp6 from *F. fulva* is known to sequester chitin oligomers to evade host defence responses in tomato (78), other LysM domain-containing effectors, such as Mg3LysM from *Zymoseptoria tritici*, have been shown to protect the hyphal cell wall from hydrolysis by plant chitinases (79). It is therefore possible that ViEcp6 plays a role in chitin protection and/or sequestration during subcuticular host colonization.

Another possibility could be that *V. inaequalis* deacetylates chitin to chitosan during host colonization to avoid activation of plant defences, as reported for other plant-associated fungal pathogens (33, 35, 80, 81) and endophytes (34). In line with this, we observed that runner hyphae and stromata developed in culture and *in planta* were completely covered with chitosan. This indicates that chitosan is the main surface-exposed carbohydrate present on these structures. Interestingly, chitosan was also observed on tubular hyphae developed on the surface of hypocotyls and chitosan production could be induced during growth on the surface of CMs coated with apple wax. These results suggest that chitosan induction is triggered by a plant-derived cue present in the apple wax. Additionally, chitosan labelling was observed at the periphery of infection-like structures developed inside CMs, indicating that another trigger, such as pressure, might be involved. It is important to note here, however, that although the OGA^488^ probe is specific for chitosan (60), binding to the positively charged groups of the carbohydrate, it cannot be ruled out that it also binds to positively charged proteins present on the surface of infection structures.

In addition to preventing activation of the plant immune system, chitosan has also been reported to be important for cell adhesion and morphogenesis in fungi (81-84). Therefore, chitosan may play other functions in *V. inaequalis*. The deacetylation of chitin to chitosan is catalysed by CDAs, and an inspection of the *V. inaequalis* genome revealed that this fungus has eight predicted CDA-encoding genes, three of which (*ViCDA1, ViCDA4, ViCDA7*) are predicted to be functional. One of these, *ViCDA1*, encodes a protein with a predicted N-terminal signal peptide, and is constitutively expressed in culture and during host colonization. Secreted CDAs are usually assumed to deacetylate chitin to chitosan to prevent activation of the plant immune system (50, 85). Therefore, it seems plausible that ViCDA1 is the main enzyme responsible for deacetylating chitin to chitosan on the surface of *V. inaequalis* infection structures. In any case, the recently developed CRISPR-Cas9 method for reverse genetics in this fungus (86) will serve to investigate the putative role of ViCDA1 and other CDAs in *V. inaequalis*.

In conclusion, we have detailed, for the first time, the cell wall carbohydrate composition of *V. inaequalis* during growth in culture. Furthermore, by assessing the expression profile of genes putatively associated with cell wall biogenesis, as well as by monitoring the localization of cell surface-associated carbohydrates using CLSM, we have also provided new insights into how this fungus differentiates and protects its infection structures during host colonization. Importantly, as the first study of its kind for a subcuticular pathogen, our research provides a foundation for understanding not only how this class of plant-pathogenic fungi causes disease, but also how they can potentially be controlled.

## Materials and methods

### *V. inaequalis* isolate

*V. inaequalis* isolate MNH120, also known as ICMP 13258 and Vi1 (77, 87), derived from a monospore culture, was used in this study.

### Preparation of fungal material for glycosidic linkage analysis and proteomics

Fungal material for glycosidic linkage analysis and proteomics was prepared by inoculating 100 µl of 10^5^–10^6^ conidia from *V. inaequalis* on CMs (Waugh Rubber Bands; Wellington, New Zealand) overlaying PDA (Scharlab, S.L; Senmanat, Spain) and culturing for 5 days at 22°C under a 16 h light/8 h dark cycle. Following culturing, fungal biomass, comprising a mix of tubular hyphae and asexual conidia was harvested from the surface of the CMs using an L-shaped spreader, washed three times by centrifugation at 4,000 g for 15 min, frozen at −20°C, freeze-dried, and then ground to a fine powder using liquid nitrogen.

### Glycosidic linkage analysis

Cell wall material from *V. inaequalis* grown in culture on the surface of CMs overlaying PDA at 5 dpi was prepared in triplicate (i.e. as three technical replicates) using a previously described protocol (88). The cell wall preparations were subjected to permethylation and GC/EI-MS analysis (88). Partially methylated alditol acetate (PMAA) derivatives were separated and analysed by gas chromatography (GC) on a SP-2380 capillary column (30 m x 0.25 mm [inner diameter]; Supelco) using an HP-6890 GC system with an HP-5973 electron-impact mass spectrometer (EI-MS) as a detector (Agilent Technologies; CA, US). The temperature was programmed to increase from 180°C to 230°C at a rate of 1.5°C/min. The mass spectra of the fragments obtained from the PMAA derivatives were compared with reference derivatives.

### Proteomic analysis

Approximately 10 mg of powdered fungal material from *V. inaequalis* grown in culture on the surface of CMs overlaying PDA at 5 dpi in triplicate (i.e. as three technical replicates) was boiled in SDS buffer (75 mM Tris-HCl buffer pH 6.8 containing 3% (w/v) SDS, 100 mM DTT, 15% (w/v) glycerol and 0.002% bromophenol blue) for 5 min at 95°C. Insoluble material was then removed by centrifugation at 14,000 *x g* for 10 min and the supernatant was loaded on a 12% Mini-Protean TGX SDS-PAGE system (Bio-Rad; CA, USA). After staining with Coomassie Blue (ThermoScientific, MA, USA), the gel lane was cut into 10 bands and the proteins were subjected to in-gel digestion with trypsin (Promega, Madison, WI, United States) as previously described (Leijon et al., 2016).

Peptide analysis was performed with reverse-phase liquid chromatography electrospray ionization–tandem mass spectrometry (LC–ESI–MS/MS) using a nanoACQUITY Ultra Performance Liquid Chromatography system coupled to a Q-TOF mass spectrometer (Xevo Q-TOF, Waters; MA, USA). The in-gel tryptic peptides were resuspended in 0.1% trifluoroacetic acid (TFA) and loaded on a C18 trap column (Symmetry 180 μm × 20 mm, 5 μm, Waters) that was then washed with 0.1% (v/v) formic acid at 10 μl/min for 5 min. The samples eluted from the trap column were separated on a C18 analytical column (75 μm × 150 mm, 1.7 μm, Waters) at 250 nl/min using 0.1% formic acid as solvent A and 0.1% formic acid in acetonitrile as solvent B in a stepwise gradient: 0–10% B (0–5 min), 10–30% B (5–70 min), 30–40% B (70–72 min), 40–90% B (72–75 min), 90% B (75–80 min), and 90–0.1% B (80–85 min). The eluting peptides were sprayed in the mass spectrometer (capillary and cone voltages set to 2.1 kV and 45 V, respectively), and MS/MS spectra were acquired using automated data-directed switching between the MS and MS/MS modes using the instrument software (MassLynx v4.0 SP4). The five most abundant signals of a survey scan (400–1300 m/z range, 1 sec scan time) were selected by charge state, and collision energy was applied accordingly for sequential MS/MS fragmentation scanning (100–1800 m/z range, 1 sec scan time).

The resulting MS raw data files were processed using Mascot Distiller (v2.4.3.2, Matrix Science, London, UK), and the resulting files were submitted to a local Mascot (Matrix Science, v2.3.1) server using the *Venturia* protein database (25,153 sequences) (48). The following settings were used for database search: trypsin-specific digestion with two missed cleavages, ethanolated cysteine as fixed and oxidized methionine as variable modifications, peptide tolerance of 200 ppm and fragment tolerance of 0.2 Da. Only those peptides with mascot scores exceeding the threshold for statistical significance (*p* <0.05) were retained. The peak list files generated from the mass spectrometry raw data have been deposited to the MassIVE database (accession number MSV000090342 (doi:10.25345/C56M3379F).

### Plant growth conditions and infection of apple material

Open-pollinated *Malus* x *domestica* cultivar ‘Royal Gala’ apple seeds (Hawke’s Bay, New Zealand) were germinated at 4°C in moist vermiculite containing 100 mg/ml Thiram fungicide (Kiwicare Corporation Limited; Christchurch, New Zealand) for approximately two months in the dark. Once germinated, apple seedlings were planted in potting mix (Daltons™ premium potting mix; Daltons, Matamata, New Zealand) and grown under a 16 h light/8 h dark cycle with a Philips SON-T AGRO 400 Sodium lamp, at 20°C with ambient humidity for 4-to-6 weeks. For the growth of etiolated hypocotyls, a fast-germination method was used. Seeds were sterilized with 5% ethanol for 5 min, rinsed five times with sterile MilliQ water, and soaked in MilliQ water overnight. The next morning, the testa of the apple seeds were peeled away with sterile forceps until the underlying white embryo was uncovered. Then, the white embryo was placed in Murashige and Skoog (MS) agar medium (2. 25 g/L MS (Sigma-Aldrich; Castle Hill, Australia), 10 g/L agar, pH 5.8 (KOH)) inside 50 ml Falcon™ tubes, with the tip of the radical submerged in the agar. Once the apple seeds had germinated and the cotyledons were fully expanded, the seedlings were transferred to potting mix soil and grown at 20°C with ambient humidity in the dark for two weeks. *V. inaequalis* infection of detached apple leaves and hypocotyls was performed as described by Rocafort et al. (2022) and Shiller et al. (2015), respectively.

### Extraction of apple wax and coating of cellophane membranes

Approximately ten fruit from apple cultivar ‘Royal Gala’, acquired from a local supermarket in New Zealand, were used for wax extraction. To extract the wax, each apple was dipped a total of five times, each for 10 sec, in 200 ml chloroform contained within a glass beaker. Following this step, the chloroform in the beaker was evaporated off at room temperature (RT) until the extracted wax had completely dried out. To coat the CMs with apple wax, a Kimwipe (Kimtech; Milsons Point, Australia) was dipped in chloroform, wiped over the surface of the wax in the beaker, and then transferred by wiping onto the surface of CMs until an homogeneous white coating was observed. Coating of the CMs with apple wax was performed under sterile conditions as much as possible, with coated CMs subsequently subjected to 15 min of UV exposure to ensure sterility.

### Confocal laser scanning microscopy

Cross-sections of detached apple leaves infected with *V. inaequalis* at 7 dpi were prepared for labelling as described previously (9). Briefly, leaf tissue was fixed in paraformaldehyde and 2.5% (v/v) glutaraldehyde in 0.1 M phosphate buffer (pH 7.2) and then embedded in LR White resin (London Resin, Reading, UK), with samples subsequently cross-sectioned from the resin-embedded material.

All the other samples (non-cross-sectioned), CMs associated with *V. inaequalis* at 6-7 dpi, as well as detached apple leaves and etiolated hypocotyls infected with *V. inaequalis* at 7 dpi, were fixed in 95% ethanol and stored at 4°C until required. Additionally, to enhance the penetrability of the carbohydrate-specific probes and antibodies used for labelling (see below), *in planta* samples were macerated with 10% KOH at RT for 4 h. For α-1,3-glucan and β-1,3-glucan labelling, non-cross section CM samples were gently scratched with sandpaper (200 grit) to generate entry points for the antibody. All samples were washed 3 times with 1 ml phosphate-buffered saline (PBS) buffer (pH 7.4) prior to labelling.

To facilitate the visualization of *V. inaequalis* infection structures in apple hypocotyls, fungal nuclei were stained with 0.002% (w/v) propidium iodide (PI) in PBS buffer (pH 7.4) containing 0.02% Tween 20. For the labelling of chitin and chitosan, carbohydrate-specific probes were used. More specifically, for WGA^488^ labelling of chitin, samples were vacuum-infiltrated in the dark for 30 min in a solution containing 0.1 mg/ml of WGA^488^ probe and 0.02% Tween 20 in PBS buffer (pH 7.4). For calcofluor white labelling of chitin, samples were incubated in 300 mM sodium hydroxide (NaOH) for 1 h prior to incubation with 0.01% calcofluor white (Sigma-Aldrich) in PBS (pH 7.4). For OGA^488^ labelling of chitosan, samples were vacuum-infiltrated in the dark for 30 min with a solution of 0.1% (w/v) OGA^488^ and 0.02% Tween 20 in PBS buffer (pH 7.4). For this purpose, a 1 mg/ml stock solution of OGA^488^ in 0.1 M sodium acetate buffer (pH 4.9) was kindly provided by Jozef Mravec from the University of Copenhagen (60). In all cases, the probe used for labelling was vacuum-infiltrated into samples for 20 min, twice, in the dark.

For the labelling of β-1,3-glucan in cross-sectioned and non-cross-sectioned samples, and α-1,3-glucan in non-cross-sectioned samples, carbohydrate-specific antibodies were used. More specifically, samples were first washed three times with 1 ml PBS buffer (pH 7.4) and blocked with 3% bovine serum albumin (BSA, Gibco; Maryland, USA) in PBS buffer for 1 h at RT in a rotatory shaker (John Morris Scientific; Palmerston North, New Zealand). Then, samples were washed with 1 ml PBS buffer three times and vacuum-infiltrated for 20 min with 0.1 mg/ml of primary mouse anti-1,3-β-glucan antibody (Biosupplies; Sydney, Australia) for β-1,3-glucan labelling or 0.1 mg/ml mouse MOPC-104E primary antibody (Sigma-Aldrich) for α-1,3-glucan labelling, and then incubated overnight with shaking (30 rpm) at 4°C. The next day, samples were washed three times with 1 ml PBS buffer (pH 7.4) and vacuum-infiltrated for 20 min with 0.1 mg/ml anti-mouse CF-488 secondary antibody (Biotum; California, USA) in PBS buffer (pH 7.4) and incubated with shaking at 30 rpm, at RT in the dark. Finally, samples were washed three times with 1ml PBS buffer.

CLSM was performed on labelled samples using a Leica SP5 DM6000B confocal microscope (488 nm argon and 405 nm UV laser) (Leica Microsystems, Manheim, Germany), with images produced using ImageJ 1.x software (NIH) (89). Here, multiple optical sections (z-stacks) were projected into a single image as maximum or average intensity projections. PI was excited at 561 nm using a DPSS laser with an emission spectrum of 561–600 nm. WGA^488^, and OGA^488^ and CF-488 were excited using a 488 nm Argon laser (power ~30%) with an emission spectrum of 498–551 nm. Calcofluor white was excited using a 405 nm UV laser with an emission spectrum of 445–455 nm. For all samples, an appropiate non-stained control was performed (**Figure S5**).

### Annotation of fungal cell wall enzymes

The predicted *V. inaequalis* isolate MNH120 protein catalogue from Rocafort et al. (2022) (10.5281/zenodo.6233646) was used in this study. Proteins that were putatively associated with fungal cell wall biogenesis were identified based on CAZyme annotation and KEGG classification, and decisions about the potential classification of enzymes involved in cell wall biosynthesis were further assessed by InterProscan scan annotation (in conjunction with the Pfam, HAMAP, MOBIDB, PIRSF, PROSITE and SUPERFAMILY tools). Here, all InterProScan and CAZyme annotations, as well as KEGG classifications, were provided by Rocafort et al. (2022). To identify CDAs, proteins with a CE4 CAZyme annotation or with ‘polysaccharide deacetylase domain’ (PF01522) were annotated as CDAs. To further investigate other putative CDAs in the genome a BLASTp (90) search was performed against the *V. inaequalis* MNH120 genome (77) using the protein sequence of CDAs from pathogens whose activity has been experimentally shown: *P. graminis* (GMQ_17027) (91), *Colletotrichum lindemuthianum* (ClCDA) (AAT68493) (50), *Pestalotiopsis* sp. (PesCDA) (KY024221) (85) and *M. oryzae* (MGG_12939, 14966, 09159, 04172, 08774, 01868, 08356, 05023, 04704 and 03461) (81).

Although the enzymatic activity of the *M. oryzae* CDAs has not been experimentally shown, their function as potential CDAs has been reported by gene deletion studies (31, 33, 81). The E-value cut-off used for all BLASTp searches was 1E-02. To investigate whether the putative CDAs had the conserved amino acid residues required for catalytic activity and metal binding, an alignment was performed between all of the predicted *V. inaequalis* CDAs and the functionally characterized CDAs from above, ClCDA and PesCDA (92), using the MUSCLE plugin of Geneious v9.0.5.

To classify CHSs, the InterProScan domain annotation from Rocafort et al. 2022 was used and an alignment of all *V. inaequalis* CHSs was performed using the MUSCLE plugin of Geneious v9.0.5 (92). The CHSs from the *Aspergillus nidulans* (93), *Neurospora crassa* (94), *Botrytis cinerea* (95) and *Ustilago maydis* were included in this alignment as a reference. To determine if the predicted CHSs of *V. inaequalis* had the conserved motifs required for CHS activity, a protein sequence alignment was performed with the functionally characterized CHS from *N. crassa* (94) as a reference.

### Gene expression analysis

Pre-existing *V. inaequalis* gene expression data from Rocafort et al. (2022) (GEO Series accession: GSE198244) were used for the gene expression analysis, and DEGs were identified using DESeq2 package v1.32.0 (96). Genes up- or down-regulated at one or more *in planta* infection time points, relative to growth in culture, with a log_2_ fold change of +/-1.5 and a *p* value of 0.01 were considered differentially expressed. Volcano plots were generated using ggplots2 v3.3.5 (97), while gene expression heatmaps were generated using Complexheatmap v2.9.1 (98).

## Funding

MRF and CHM were supported by the Marsden Fund Council from Government funding (project ID 17-MAU-100), managed by Royal Society Te Apārangi.

## Author Contributions Statement

CM, MR, JB, VB, MA and KP conceived the project. MR, VS, SM and PS conducted the experiments. MR performed the bioinformatic analyses. MR, CM, VS, VB, JB, KP and RB provided critical input in experimental design or data analysis. MR, CM, VS and VB wrote the manuscript. All authors read, revised, and approved the final manuscript.

## Conflicts of Interest Statement

The authors declare no conflicts of interest.

## Notes

### Competing Interest Statement

The authors have declared no competing interest.

