## Supplementary Information for "Cell wall carbohydrate dynamics during the differentiation of infection structures by the apple scab fungus, *Venturia inaequalis*"

**Supplementary file 1: *Venturia inaequalis* genes that encode enzymes putatively associated with fungal cell wall biogenesis.** Enzymes were classified according to their predicted roles in cell wall biogenesis, as based on predicted 'Kyoto Encyclopedia of Genes and Genomes' (KEGG) and InterProScan annotations. CBM, carbohydrate-binding module; CDA, chitin deacetylase; CE, carbohydrate esterase; CHS, chitin synthase; GH, glycoside hydrolase; GT, glycosyl transferase; PL, polysaccharide lyase. * Only three CE4 proteins were predicted to be a carbohydrate-active enzyme (CAZYme) using dbCan2.

**Table S1. *Venturia inaequalis* carbohydrate-active enzymes (CAZymes) putatively involved in fungal cell wall biogenesis and modification.** Enzymes were classified according to their predicted roles in cell wall biogenesis, as based on predicted 'Kyoto Encyclopedia of Genes and Genomes' (KEGG) and InterProScan annotations. CBM, carbohydrate-binding molecule; CE, carbohydrate esterase; GH, glycoside hydrolase; GPI, glycosylphosphatidylinositol; GT, glycosyl transferase; PL, polysaccharide lyase; UDP, uridine diphosphate. * Only three CE4 proteins were predicted to be a carbohydrate-active enzyme (CAZYme) using dbCan2.

| Enzyme | CAZyme family | Copy number |
|---|---|---|
| **CHITIN METABOLISM** | | |
| Chitin synthase | GT2 | 8 |
| Chitinase | GH18 | 5 |
| Hexosaminadase | GH20 | 1 |
| β-N-acetylhexosaminidase | GH3 | 1 |
| Chitin deacetylase | CE4 | 8* |
| **GLUCAN METABOLISM** | | |
| β-1,3-glucan synthase | GT48 | 1 |
| α-1,3-glucan synthase | GH13+GT15 | 2 |
| 1,3-β-glucanosyltransferase | GH72 | 6 |
| β-glycanase | GH16 | 13 |
| 1,3-β-glucanase | GH128, GH152, GH64, GH81 | 8 |
| β-glucosidase | GH1, GH132, GH17, GH3, GH5, GH55 | 29 |
| 1,4-α-glucan branching enzyme | CBM48+GH13_8 | 1 |
| α-amylase | GH13 | 7 |
| alpha-1,6-glucohydrolase | GH13 | 2 |
| glucan 1,4-alpha-glucosidase | GH15 | 2 |
| α-glucosidase | GH31, GH71 | 6 |

| | | |
|---|---|---|
| **MANNOSE METABOLISM** | | |
| α-1,2-mannosyltransferase | GT15, GT4, GT71 | 7 |
| α-1,3-mannosyltransferase | GT69 | 7 |
| α-1,6-mannosyltransferase | GT22, GT34 | 6 |
| α-1,3/α-1,6-mannosyltransferase | GT4 | 1 |
| GPI-mannosyltransferase | GT22, GT50, GT76 | 4 |
| β-1,4-mannosyltransferase | GT33 | 1 |
| Dolichyl-phosphate-mannose-protein mannosyltransferase | GT39 | 3 |
| Mannan polymerase complex MNN9/ANP1 | GT62 | 2 |
| α-1,2-mannosidase | GH92, GH47 | 13 |
| Mannan endo-1,6-α-mannosidase | GH76 | 8 |
| Mannosyl-oligosaccharide glucosidase | GH63, GH31 | 3 |
| α-mannosidase | GH38 | 1 |
| β-mannosidase | GH2 | 1 |
| Mannosyl-glycoprotein endo- β -N-acetylglucosaminidase | GH85 | 1 |

| | | |
|---|---|---|
| **GALACTOSE METABOLISM** | | |
| Inositol 3-alpha-Galactosyltransferase | GT8 | 1 |
| α-galactosidase | GH27 | 6 |
| β-galactosidase | GH2 | 1 |
| α-1,4-galactosaminogalactan hydrolase | GH135 | 4 |
| α-1,4-polygalactosaminidase | GH114 | 9 |
| N-acetylgalactosamine deacetylase | CE18+CBM87 | 4 |
| Endo- β -1,6-galactanase | GH5 | 1 |
| UDP-glucose 4-epimerase | - | 3 |

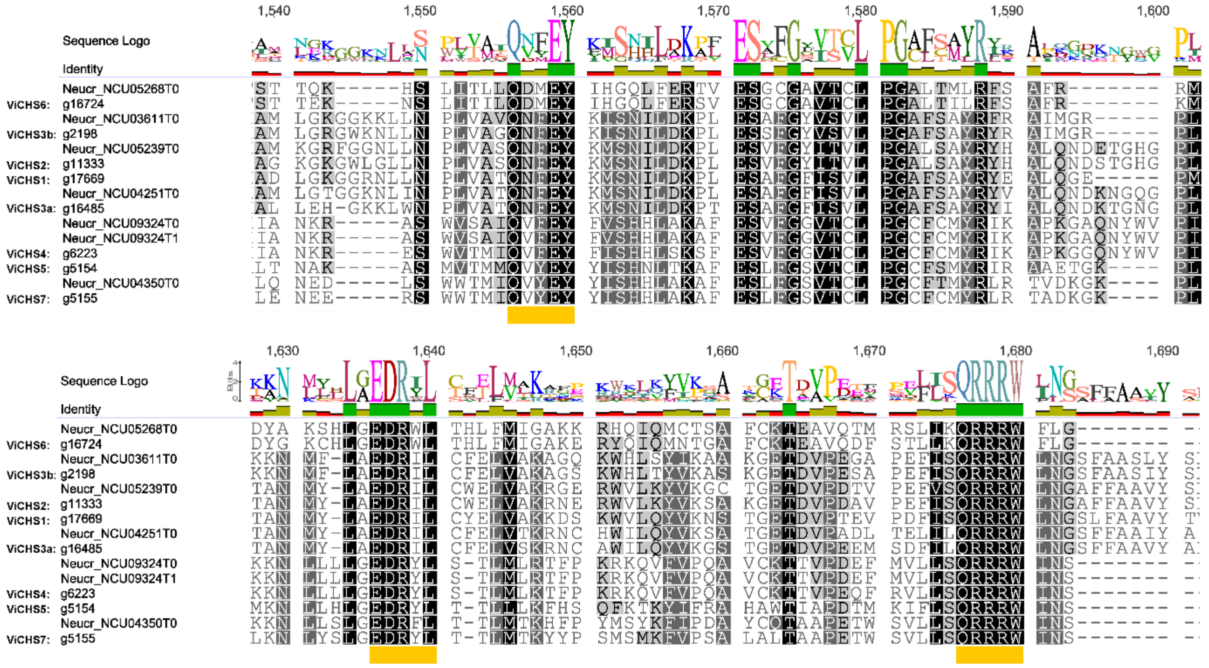

**Figure S1.** Multiple sequence alignment of putative chitin synthase (CHS) proteins from *Venturia inaequalis*. The CHS proteins from *Neurospora crassa* (Neucr) were used as reference for alignment. Alignment generated using the MUSCLE plugin of Geneious v9.0.5 in conjunction with full-length protein sequences. The conserved motifs for catalytic activity are highlighted under the alignment with orange boxes. Amino acids are coloured based on similarity, with the most similar amino acids coloured black.

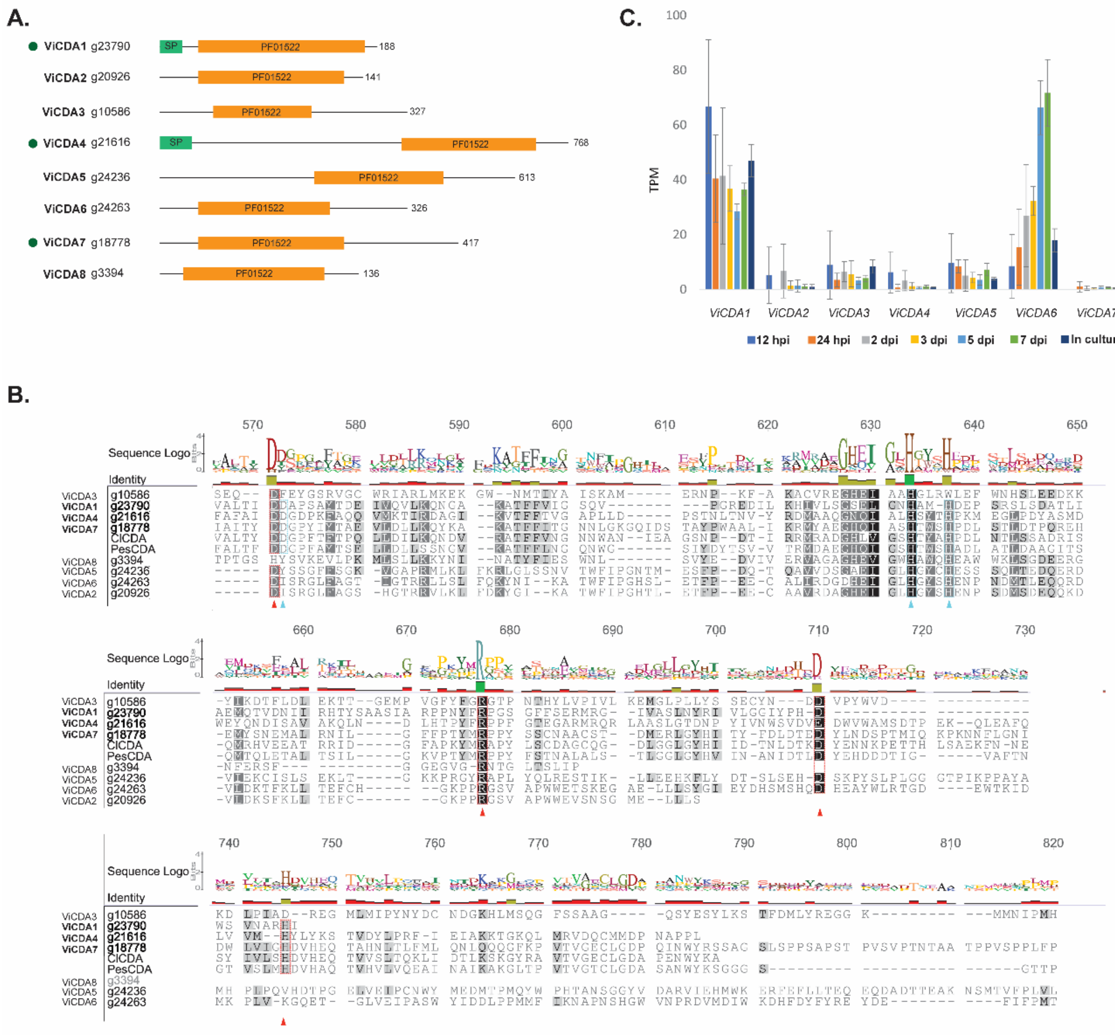


**Figure S2. Predicted chitin deacetylases (CDAs) of *Venturia inaequalis*. A.** Domain organization of putative CDA proteins from *V. inaequalis*. Only ViCDA1, ViCDA4 and ViCDA7 display all conserved amino acid residues required for catalytic activity and metal binding and, therefore, are likely to be active (labelled with green circles). Protein lengths are shown. SP, Signal peptide; PF01522, Protein family (Pfam) polysaccharide deacetylase domain. **B.** Multiple sequence alignment of putative CDA proteins from *V. inaequalis*. CDA proteins from *Colletotrichum lindemuthianum* (ClCDA) and *Pestalotiopsis* sp. (PesCDA) were used as references for alignment. Alignment generated using the MUSCLE plugin of Geneious v9.0.5 in conjunction with full-length protein sequences. The conserved amino acids for catalytic activity are highlighted with red triangles under the alignment and with dashed red boxes. Conserved amino acids for metal binding are highlighted with cyan triangles under the alignment and with dashed cyan boxes. Amino acids are coloured based on similarity, with the most similar amino acids coloured black. **C.** Expression of the eight predicted *V. inaequalis CDA* genes during growth of the fungus *in planta* (12 hours post-inoculation [hpi] to 7 days post-inoculation [dpi]) and in culture on the surface of cellophane membranes (CMs) (7 dpi). Gene expression is shown as transcripts per million (TPM). Error bars represent standard deviation across four biological replicates.

**Supplementary file 2:** List of genes encoding putative cell wall biogenesis proteins from *Venturia inaequalis* that are up- or down-regulated at one or more *in planta* time points during infection of apple leaves (12 hours post-inoculation [hpi] to 7 days post-inoculation [dpi]), relative to growth of the fungus in culture on the surface of cellophane membranes (CMs) overlaying potato dextrose agar (7 dpi).

**Supplementary file 3:** List of proteins identified by mass spectrometry from *Venturia inaequalis* grown on the surface of cellophane membranes overlaying potato dextrose agar at 5 days post-inoculation, and short-list of proteins putatively associated with fungal cell wall biogenesis from this list. Only peptides exceeding the threshold for statistical significance ($p < 0.05$) were selected.

**Supplementary file 4**: Peptide coverage of proteins identified by mass spectrometry from *Venturia inaequalis* grown on the surface of cellophane membranes overlaying potato dextrose agar at 5 days post-inoculation that are putatively associated with fungal cell wall biogenesis.

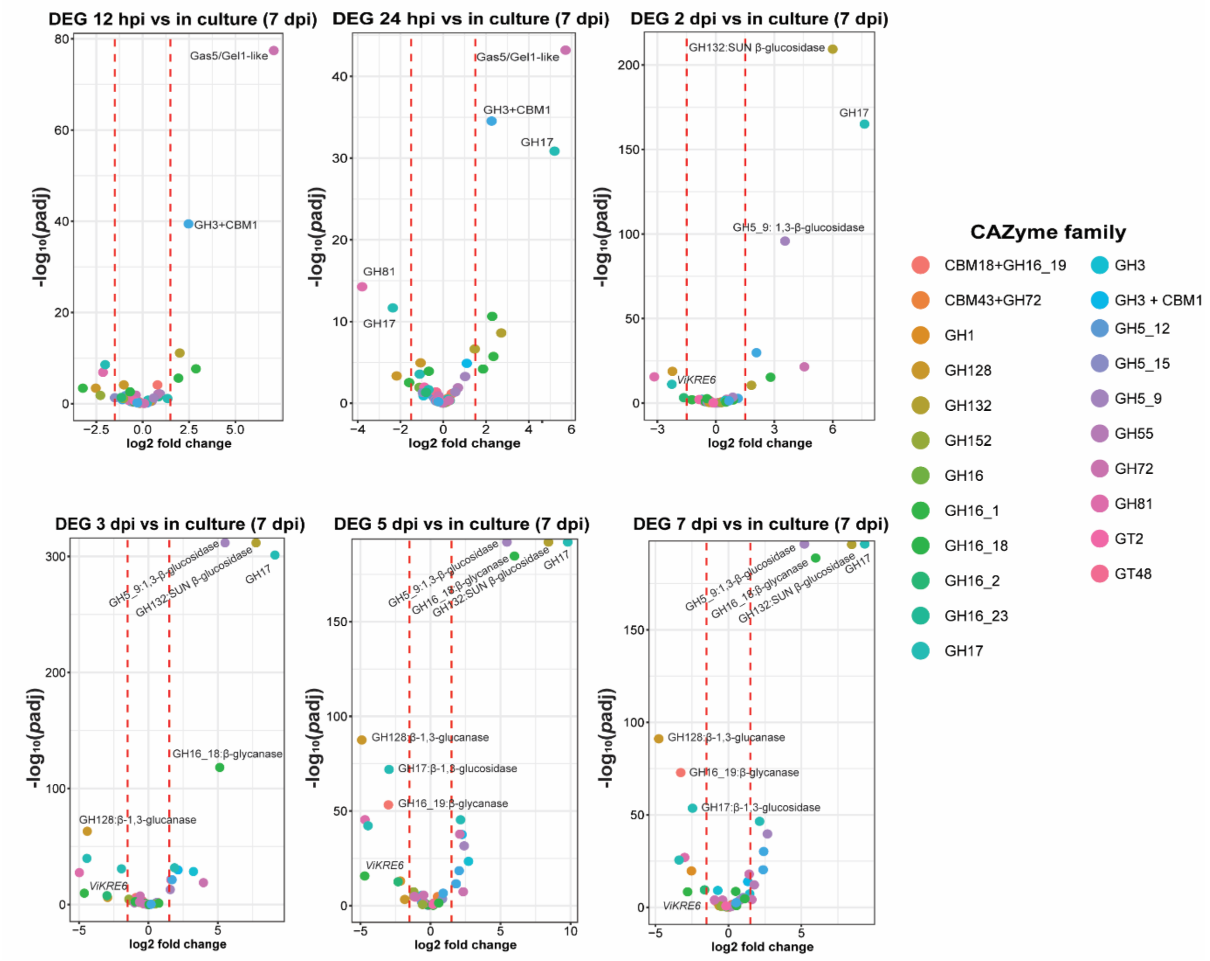


**Figure S3. Volcano plots illustrating genes of *Venturia inaequalis* that are both putatively associated with β-glucan metabolism and are up- or down-regulated during one or more *in planta* time points compared with growth in culture on the surface of cellophane membranes (CMs) overlaying potato dextrose agar.** *In planta* time points were at 12 and 24 hours post-inoculation (hpi), as well as 2, 3, 5 and 7 days post-inoculation (dpi), while growth in culture was at 7 dpi. Dashed red lines indicate the 1.5 $\log_2$ fold change used in this study to identify significant (*p*-adjusted, *p*adj) differentially expressed genes (DEGs). Each dot represents one gene, coloured by carbohydrate-active enzyme (CAZyme) classification. Only genes with a minimum expression of 10 transcripts per million (TPM) at one or more *in planta* time point are shown. CBM, carbohydrate-binding module; GH, glycoside hydrolase; GT, glycosyl transferase.

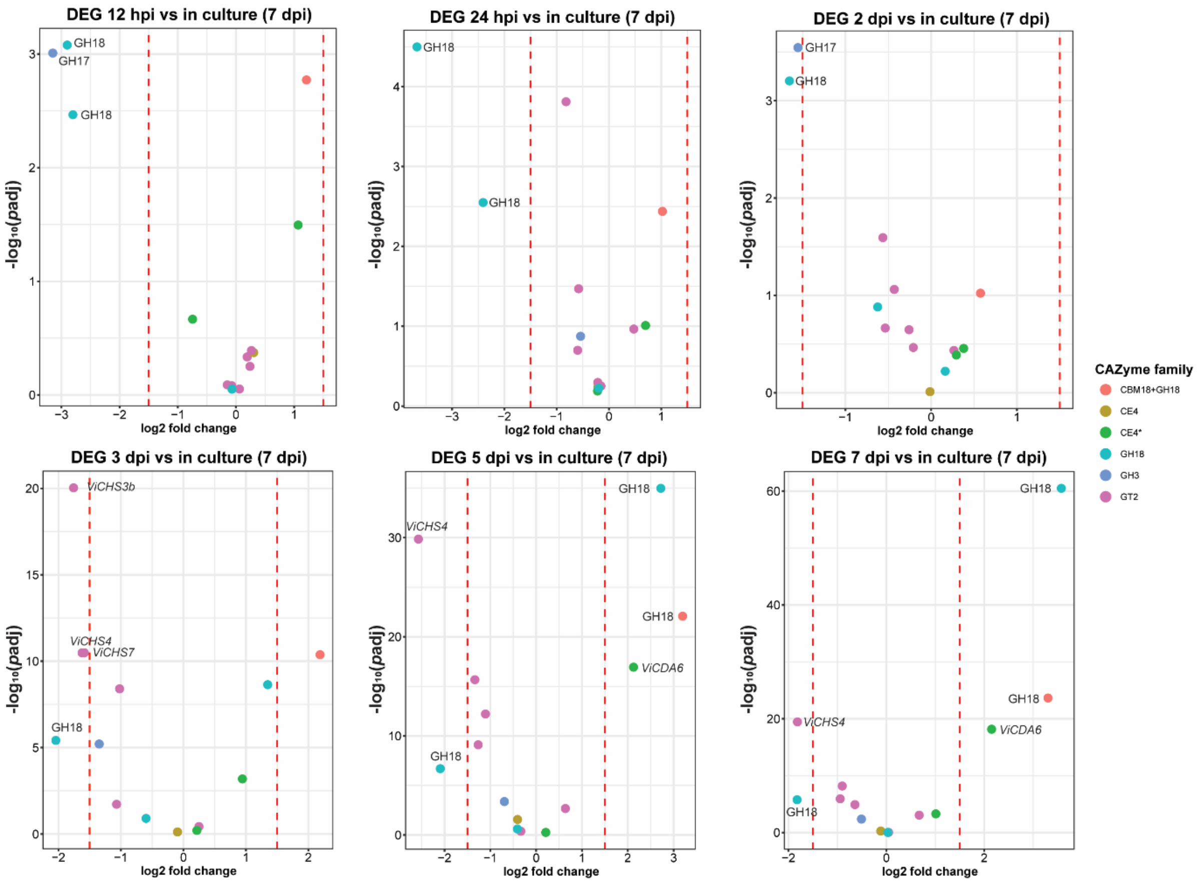


**Figure S4. Volcano plots illustrating genes of *Venturia inaequalis* that are both putatively associated with chitin (N-acetylglucosamine) metabolism and are up- or down-regulated during one or more *in planta* time points compared with growth in culture on the surface of cellophane membranes CMs overlaying potato dextrose agar.** *In planta* time points were at 12 and 24 hours post-inoculation (hpi), as well as 2, 3, 5 and 7 days post-inoculation (dpi), while growth in culture was at 7 dpi. Dashed red lines indicate the 1.5 $\log_2$ fold change used in this study to identify significant (*p*-adjusted, *padj*) differentially expressed genes (DEGs). Each dot represents one gene, coloured by carbohydrate-active enzyme (CAZyme) classification. Only genes with a minimum expression of 10 transcripts per million (TPM) at one or more *in planta* time point are shown. CBM, carbohydrate-binding module; CE, carbohydrate esterase; GH, glycoside hydrolase; GT, glycosyl transferase. Asterisk (*) indicates CE family 4 (CE4) proteins were only predicted using a protein family (pfam) domain search.

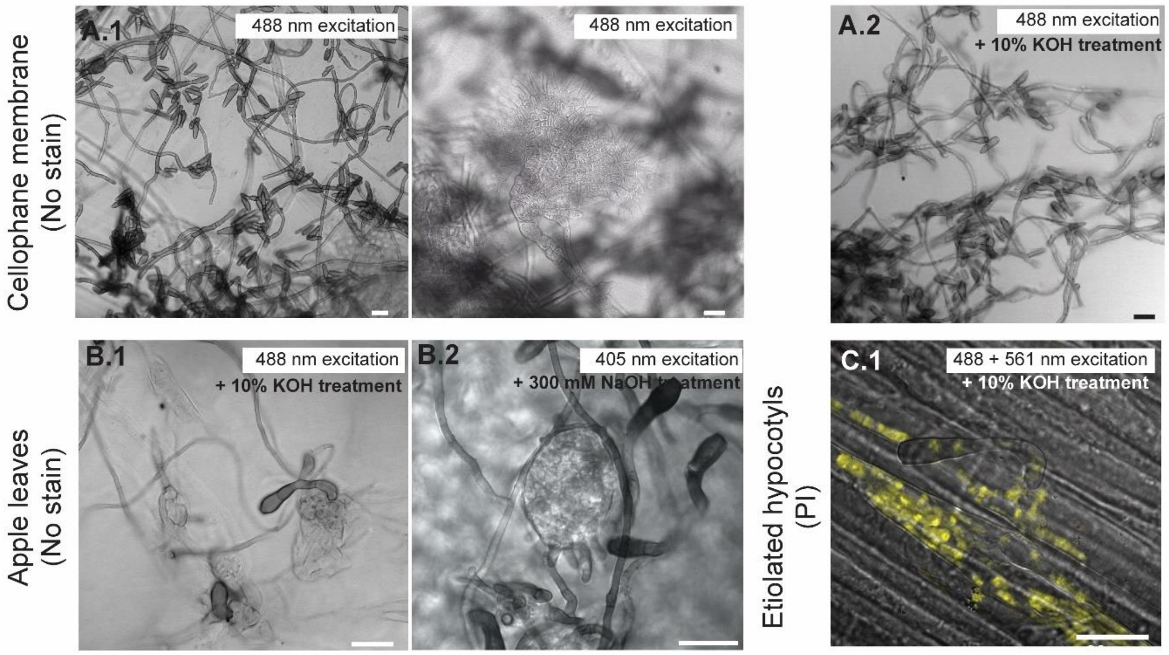


**Figure S5: Representative negative controls to ensure that the emission signals observed during confocal laser scanning microscopy (CLSM) were derived from specific labelling and not autofluorescence**. Samples of non-labelled cellophane membranes (CMs) (**A. 1**) associated with *V. inaequalis* or treated with 10% KOH (**A. 2**) did not emit any signal when excited at 488 or 405 nm by CLSM. Likewise, non-labelled *V. inaequalis*-infected apple leaf (**B.1**) treated with 10% KOH or *V. inaequalis*-infected apple leaf treated with 300 mM NaOH (**B. 2**) did not emit any autofluorescence signal. Finally, etiolated apple hypocotyls treated with 10% KOH (**C. 1**) were only stained with propidium iodide (PI) to label fungal nuclei, and only PI-derived signal was observed after excitation at 561 nm. No emission was observed after excitation at 488 nm. All scale bars: 20 µm.

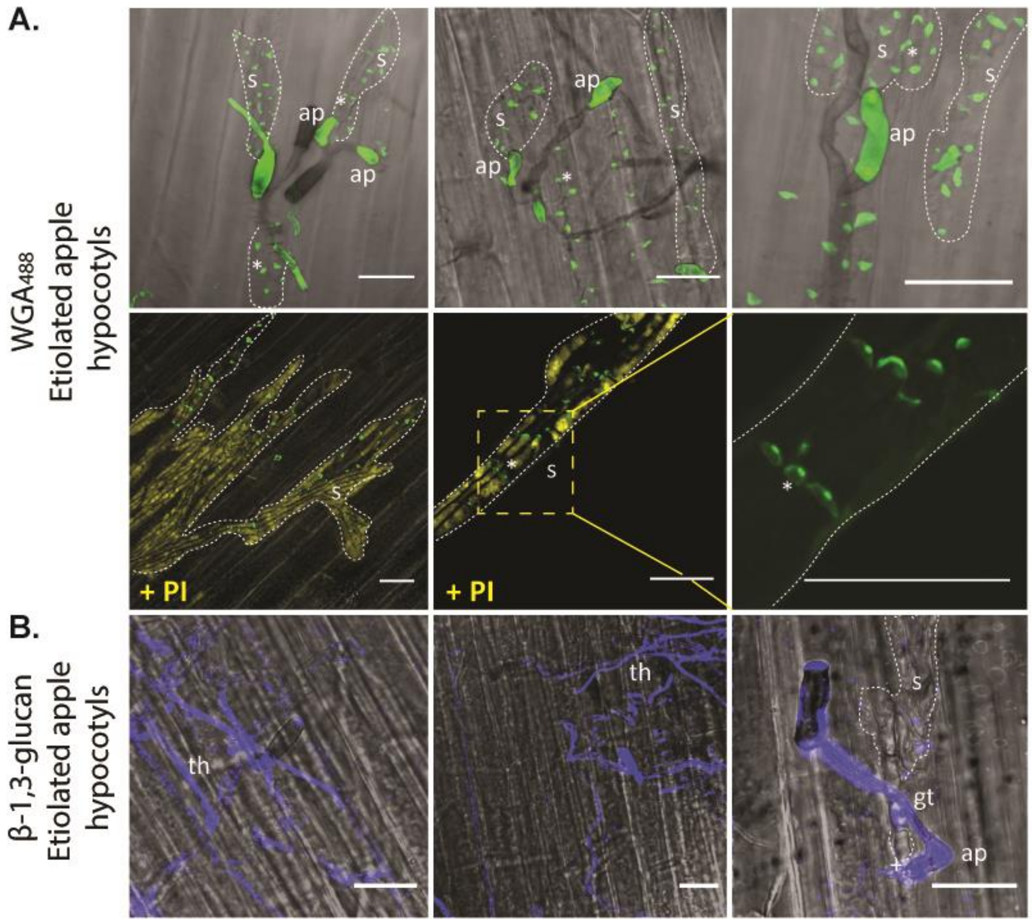


**Figure S6**. **Label-accessible chitin and β-1,3-glucan on the surface of infection structures developed by *Venturia inaequalis* in and on etiolated apple hypocotyls. A.** The fluorophore-labelled lectin $WGA_{488}$ was used to visualize chitin (green), and propidium iodide (PI, yellow) was used to stain fungal nuclei, in conjunction with confocal laser scanning microscopy. Dashed yellow squares indicate zoomed-in areas. **B.** The monoclonal anti-β-1,3-glucan primary antibody and CF-488 secondary antibody were used to label β-1,3-glucan (blue). All scale bars: 20 µm. ap, appressorium; s, stroma; th, tubular hyphae; *: septa; dashed white lines highlight stromata.
