## Supplementary File 4 for "Cell wall carbohydrate dynamics during the differentiation of infection structures by the apple scab fungus, *Venturia inaequalis*"

**Peptide detected**

Modification: X, oxidation; X, ethanolyl

### Glucose metabolism: α-glucan

>g23025: CBM48+GH13_8

MQDAMASTSATVGDGKSNIPNDGTGVVTLDPWLEPFKDSLRSRFAKAQSWIK**TINDTEGGLEK**FSRGY
EK**FGFNVLPNNDVVYR**EWAPNALR**AYLIGDFNNWNR**DSHQMTKDEYGAFDITIPAVNGQPAIPHDSKI
KISLVIPNDGHRAER**IPAWITRVTQDLNVSPVYDARFWNPPANEK**YIFKNKPPPQPKSVRIYEAHVGI
SSWEGKVATYKEFTRNVLPRIKNDGYNVIQLMAIMEHAYYASFGYQINSFFAASSRYGLPDELKELVD
TAHGMGITVLLDVVHSHASKNVLDGLNTFDGSDHQYFHEGAKGRHELWDSRLFNYGNHEVLR**FLLSNL**
**RFWMEEYQFDGFR**FDGVTSMMYTHHGIGTGFSGGYHEYFGSNVDEEAVAYMMIANELLHTLYPRVITV
AEDVSGMPALCVELALGGIGFDYR**LAMAIPDLYIK**WLKEKEDIEWDMGALCFTLTNRRHGEKTIAYAE
SHDQALVGDKSLMMWLCDAELYTNMSILSVETPVITRGLSLHKMIRLITHGLGGEGYLNFEGNEFGHP
EWLDFPREGNGNSFHYARR**QFNLADDQLLR**YRFLNDFDSKMQWTEEKYGWLHSPQAYVSLKNETDKVI
VFERAGLLWIFNFHPNQSYTDYR**VGVEEAGTYRVVINTDSK**SFGGHGNIKEETRFFTTDFAWNGRK**NF**
**VQVYIPSR**TAMVLALEGTL

>g5073: GH31

MPNMNDYKFPSEPLANPDSTITGKNYRFTLIDDKLLRYEWSADGKFEDRASTFAINRKFPKPEFRIEE
TDSQLEIFTPRYHLIYDKQRFSENGLVVQFTSKQSEWGGEWRYGYGPQQNLGGTARTLDGVDGRCDMG
AGILSRVGYAALDDTGSMLFDGEGFVGTRLEGDRIDGYLFAYGFDLRGAMKSFYAVSGSQPVVPRWCL
GNWWSRYHDYTADEYLELMDDFKKKEIPMSVAVIDMDWHIVHGDDVPHVGWTGYTWDKKLFPDPGAFT
KALHDRKLKTTLNDHPHAGVWHHEDSYEEMAKVLGHDTTNKKPILFDPTSQEFMHAFLHVLHRNLEKE
GCDFWWIDWQQGSHSRVPGLDPLWLLNHFHFIDQEQVKGKSEALIFSRYAGPGSHRYPVGFSGDAFAT
WPSLEFQPEFTATASNIGYGWWSHDIGGHLPGFRDDECATR**WIQLGVFSPILR**LHSTRSRWMSKEPWL
YRSECETAMRDAMQLRHRLVPYIYSENANTPTSSMPLVQPIYWNFPERNIAYKFPNQFYFGSELVVSP
VVTPRDPRTNLAKTKVWVPPARHVDLLAGLIYDGDREIDVYRSLNDVPVLAK**EGTIIPLDGAK**VPANG
CGNPDAFEVLVVVGQDGNYNILEDTRDDAKPVETESPRSILIDYKQAEGKLSVDGNGRAWTFR**FISFT**
**DDLSNIK**VSSSDAEVSVTSTPTPSLTVTIPTTTDKITIELGSNPQLSVLDNTKSISDILVNYQININI
KDEIWKALTAEQATITKMGRLLSLGLEEALVGPLVEFLVADSRSINGGMTKEKQYAGVIAQGD

>g748: ViAGS1 GH13_22+GT5

MHVFVWVVLTLCARVFGAAYEASEVDFNLNQNQSATHPLDYYGEWSNHTYNKSPSNWR**FPFYTLFLDR**
LANGDPSNDNANGTLFEQDILSNQFRHGGDVAGLLDSLDYIQGMGMKGLYIAGSPFMNQPWTADSYSP
VDLTILDHHFGTIDTWRTVITAIHARGMYVVLDNTFATMGDLFGFDGYLNTTTPFSLAEHKSQWKGQR
RYLDFDIGEEYNTTCAYPRFWNETGYRIDSIYTDKMKGCYNSEFDQYGDTEAFGVYPDFQRQLTKFAS

VQDRLREWHPPVLEKINHFSCMVIKMLDIDGFRFDKATQITVDAQAGFGQYMRNCAKSVGKDNFFMPG

EITGGNTFGSIYLGRGRQPDMLPENATVATKMTNNSNSKYFIRDQGMGALDAAAFHYTIYRSLTRFLG

MDGNLEAGYDANGGTWTEAWNTMLLTNDMVNENSGIFDPRHMFGVTNQDVFRWPAISQGTERMILGMF

VTAIHLPGIPLVTFGEEQAFYVLDSTADNYIYGRQPMTASIAWQMHGCYSLGSSQYYQMPLNDSRNGC

KDDTVSYDHRDPSHPVRNILKHMYFLREQFPVLNDGWYLQDLSNQTHDVVLPGSSGKPTETGIWSVVR

KEFTGSQTLSSNQTIWFVYHNVNETTTYTFDCNSDKTSFLSAFDKDTTAKNLMYPFDEITLGSTTTTK

ANTGCVSNITMKPYEFKVYVPK**AAFVTALPMITK**FLPGHDVRVKSSVSPGEQETVDVEIQFSAVMSCD

AVTSSILLNSTVEDLRIAQIKSGSVKCSTLNSTTTRYVGDLPSAWTWKATLENVSNGIHAITVKNASR

TDGTATNSVDRFIFRVGQDDNPMVFPHTANYTRALIHTNGSSKDLYISHKAAGADKWRYSLTWGSSWS

NWENYYGGNSTLQQQNWTGTSLQKWDGTHIMAQYWSRLSGSSAVIQHADLGREGQPPRRFPHLFAQGP

YNQYGFDSGLKNKVELTKDGSWDWHFMSEWPDTVQLNVWGMNPDGQPDAAFVYGDIDADGILDRLPPS

SLSNLAINVTNGPPSPWLAWKLSIDDGTLRYTLLPVGNRWLQLALFMLMAFIPLITACLAVWAYMRGF

YEVKFNEVGVSEK**TGFMALPLALR**RGRGMKKLAQFASEDNLTNMDSAIPFSTAVAGLSEKRRTVLIAT

LEYDIDDWNIKIKIGGLGVMAQLMGKNLEHQDLIWVVPCVGGVDYPPATAAEPMEVIILGEKYDIQVQ

VHQLR**NITYVLLDAPIFR**AQSKTEPYPPRMDDLDSAVYYSAWNQCIARAIQRYPVDLYHINDYHGACA

PLYLLPETIPCCLSLHNAEFQGLWPMRTPKERAEVCSVFNLSEKTVAQYVQFGSVFNLLHAGASYLRV

HQKGVGAVGVSNKYGKRSWARYPIFWGLKAIGKLPNPDPTDTEAWDKQLPKEGDIHVDPTYEAGRGEL

KRQAQEWAGLEQRADAELFVFVGRWSVQKGVDLIADVFPAVLEHHEKVQLICIGPVIDLYGKFAALKL

GKMMEKYPGRVYSKPEFTALPPYIFSGAEFALIPSRDEPFGLVAVEFGRKGALGVGARVGGLGQMPGW

WYTVESTTSAHLLKQFKESIDTALASSQETRAMMRARSAKQRFPVQQWKEDLGILQSKCIKLHDQEVE

KHGGRLTRHHDSGAPAPPIVTLENNELRSVSSPTTGSAVEGPSSGGLKRTLSLGVRQGPGHRASGDGD

GENHPENIPEEEEYIVEHGDQTAGEEGNPDSLPLPPWPIRGPGSRFSTASAMSAVSFDPQRASTATEH

PSPVPSPAFLRPDRDGAESPGYYEPDSLVPPSPGFYQQPDANRSSLLSLNLVVGEKKDFKLQKVDPFF

TDSNMEYFHAFESKLGDLNGKNSEHDLCIEEYLIKSERTWFGKFRDARLGRTTEKGIAGSRVTSRAPS

PSPSTYRFPNDSYSEEDENDKTPLRDEFLLGDDYKPPTGLKKHAQMKVGDWPIYTLLLAIGQIISANS

YQVTLLTGSVGQAASKLYVVASIYLVASIGWWVLFRRVKSVYCLSSPFIFYGLAFLLLGMAPFVSNTT

GRGWLQNVATGIYAVASASGSLFFAMNFGDEGGAPVKDWIYRACIIQGTQQLYNAALWYWGTTFTATS

SSSGSTVASSYNAFVNSWRMTAITVPVAILMWVVGYLILVGLPSFYRQAPGAVPSFYSALARRKIVVW

TLVTVLIQNFFLSAPYGRNWQFLWASRAAPKWAVALLMVLFFLVIWLGILLVFAKLSKSHSWIVPIFA

IGLGAPRWCQMLWSTSNIGLHLPWAGGAVSSALVSRALWLWLGLLDTIQGVGFGMIFLQTLTRIHVAF

TLIAAQVLGSIATIIARAAAPNNLGPGDVFPDFSQGANHVLSKAWFWIALLFQLLVCAGYLTFFRKEQ

LNKP

>g7759: GH31

MARSSLRKVIGVISTGLFLNNAQAQGVTPTPTTATELTATIAGSVVTYSPKFTIPAAADDGANLLPNI

HDPLAVDAQTVCPGYTASNYARNAHGFSASLKLAGEACNVYGTDVDELNLNVQFQNADRLSVQITPTN

VDASNSSWYVLPETLVPRPGLDEDAKSTTIDNDFQITWSNDPTFSFTVLRRSSGDVVFSTAGSKLVYE

DQFIEFVTTMPENYNIYGLGEQIHGFRLGNNYTATIFAADAGDPIDGNIYGSHPFYLDTRYYEVDPET
GNKTLVTSNDTSASSTYESVSHGVFLRNAHAQEVLMRADNITWRTLGGSIDLFFFDGPTQPEVTKQYQ
VGAIGLPAMQQYFSFGYHQCRWGYQNWTILQEVVDNFRNYNIPLENIWTDIDYMNRYRDFDMDPNTFS
VSEGKQFLDKIHANHQHYIPIVDSAIYIPNPDNATDAYAVYDRGNSSDVFMKNPDGSEYIGAVWPGYT
VYPDWHADESVPWWVNEMVMWYKDINFDGIWIDMSEVSSFCIGSCGTGNLSLNPAHVPFLLPGEPGNL
QLEYPEGFNITNATEASSVAALSSSQAAALSSTAVSTSTESAVTSTSSASTSASYVRTTPTPGVRNVE
LPPYVINNINGQLGTHALAPNATHADGVQEYDVHNLFGHQILNATYQALLSVFPGKRPFIIGRSTFAV
SGKWAGHWGGDNNSKWPWMGFSIPQALHFGLFGIPMFGVDTCGFNGNSDEELCNRWMQLSAFFPFYRN
HNVIGTISQEPYIWGSVAEATRKAIAIRYALLPYMYTLFHSAHTTGTTVMR**ALAWEFPNDPSLANADR**
QFLLGPSILVTPVLAQGATSVNGVFPGAGKGEVWYDWYTHSAIHASAGENITIAAPLGHIPVYIRGGS
VLPLQQPGYTTFESRQNPWALLVALNLDGAATGSLYLDDGESIMQNSTLFVEFTASAGKLSVSATGEF
VDANSLANITILGLSRAPVDVALDGVLIGSGVSFDGGSGVLSVTGLAGATGGGAWSRDWVLSWS

>g8055

MFVRHRFQELIAFVGLLCVFSSAQDNCTTSIVSTAPPANGTGVQLQQFSYCGGDLNITAYIEDVNYEK
IVIVSYSDRSNKSTPVNSIALDYVSAVTGNERWQLWSAKTPIYIDGITELLKITYKAVNVNKVYTQIL
NIPVVASGRPAPAPLAAPVPYATPSGFSDDITSWLAVGALSQIATCKTRMFDNINSVGAANGTVVAST
STANPDYHYNWVRDAALTMDVVVDLYEAATVPPAVSYYENILFQYSQAR**AGQQQIGGLGEPK**FYLNNT
LFTGPWGRPQNDGPAEAAVALIDFASAYLAKNGSLEKVRKEIYDSTTYPTFAPVKRDLLYVAANWSLE
SFDLWEEESSYHFFNSLVSHRALTIGSAFAAKLNDSSTASTLSTAADAVVAGMGRFWDENRQLILYEY
GPVLKNKTSFKDIAVILGVLHGYADDDIFSYTNDQVLVSAYQIATSFLSVYPIAKVTLDSAGGVLGIP
IGRYPEDVYNGTGTAPEGGNPWYLATSTLAEFLYKATHSFTNQSSIKVSNTSLPFWTYFAPSASLAAG
TTYPSTSSPFKTAIGALEGWADAFMRRVKFHVPADGRLAEEYHRRDGIATGAEDLTWSYASVVTAAMA
RAVVRGDVGYARGLANLGFT

### Glucose metabolism: β-glucan

>g12448: GH3+CBM1

MLALAYLLLSVTSLRLVHADLELKWSYGRSPPVYPSPSGAGTGDWNDAYAKARSALAKMSNAEKANLT
IGLTGFTGCSGTSGGAASIAFPGLCLQDGPSGVRSTDLVNAYPAQLSIGASWNRTLANGVATYMGAEF
KRK**GANVALGPTIGPLGR**VALGGRNWEGYGSDPFLSGVLSAEAVLGLQKSVMACVKHLVGNEQETNRN
PSIISFQQSVSSNIDDRTMHELYLWPFQDAVKAGAASVMCSYQKINGSYGCQNSKVLNGLLKTELGFQ
GFVVSDWGAQHAGIAGAAAGLDMVMPSASFWASNALVTAVQNGSLPQTR**LDDMATR**VLASYYRLGMDS
PSYPALGIGIPAVVTAAHPLVEGRDPASKSTLLQGAVEGHVLVKNVNNTLPLKAPTLLSLFGYDAYAP
LVVNPSSSAVDRWTHGVESVTANDVQLLLIAAGLSGTAIGAATSGTLTKGGGSGSSYGPYISAPYNAF
EQQAHKDGTYLFWDFQNQNPDVSAASNACIVFINEFATESQDRQSLADVDSDKLVNNVAKKCSNTIVV
IHNAGIRLVDAWIDNVNITAVIFAHTPGQDSGRALVEVMYGKQGFSGRLPYTVGRKQSDYGSLLSPSL

PGPILSDTYLYPQSDFTEGLNIDYRDFIARNVSPRYPFGYGLTYTTFSYSNFSITPMTNNLTVPLPPA
PSAQGGNPNLFNTIARVDCTVTNTGTVEGAEVAQLYIGIPNSPPKQLRGFEKKSLQPGERKTFSFPLA
RRDLSIWSTTRQEWVLQSGNYQIYVGASVLDIKLQGVLTI

>g15895: GH1
MALVILFSLAAITLATPQSPLEFPNPAGYEFKKYSAPSLDPLWAKIASPIAPPKYTSTVVPTPEPATY
TQPNEFHPLVASHDTNLTNLKLPKNFIWGVASSAYQIEGAAKLEGKGPSIWDALSHNVPNFVADNSTG
DNVAEQYMMYKVDIARMK**GLGIPAFSPSFAWPR**FFPTGKAKDGANEEAVKHYDDVITELVSAGIKPVI
TLFHWDTPLALFGEYGAWLSPKAMDDFVDYAKFVIQRYDSVVSTWYTFNEPQYCNWQFSEYPLDGRFY
PIGGQDLSKFVGKEGKLRARFLCGHYTLLAHARVAKWYHNEFKGKGRITFKNSGNFQEPRTQSAADLR
AAQRGFDFSIGWFGGPWTDGDYPTSLRETLGDLLPTFTTEEKSLIKGSCDFYAIDPYTSFTLYSEPDA
ENCYTNRTNSGYPECTSSTQTGANGFPMGPSSDNGVSWLKSTPYGIRKFLKKIMVLFPSVPDIVVSEF
GFAEPFESRLTKMEDILWDLRRADYYQNYLDAILQSIHYDHINITGAWGWSVYDNFEWLVGSDVRFGL
QYLNYTSLERTPK**ASMFQFLNWFK**QHSA

>g16312: GT48
MSGHPPPPQGGYHDEAYDANGQPYYNDGQGYYDQNQQYEGQPHGAPVAGQDPYYDDQGYYNDNHQGGY
AQDGYYDQNGYQGDEYYDNQYYDQAQGGQPPQGYGYDGGQRRQRRGDSEEDSETFSDFTMRSDVHRAA
DMDFYGRGDERYNSYNGEQQGRGFRPPSSQISYGGNRSSGASTPVYGTEFGALPAGQRSREPYPAWTQ
DAQIPITKEEIEDVFMDLREKFGFQGDSVRNMYDHFMTLLDSRASRMTPNQALLSLHADYIGGENANY
RRWYFAAHLDLDDAVGFANMELGKGNRRTRKARRAAKKKAAENPADEAKTLEALEGDNSLEAAEYRWK
TRMNRMSQHDRVRQIALYLLCWGEANQVRFTPELLCFIFKCADDWLNSPAAQSGHFVVEEGTYLNTVV
TPLYQYMRDQGYEIQDGKYMRRERDHAQIIGYDDINQLFWYPEGIERIVMEDKTRIVDFPPAERYAKL
KEVAWKKVFFKTYKETRSWFHLIVNFNRIWVIHVTAFWFYTAYNSPTLYTKDYQQERNQKPNPPAQWS
AVALGGTLACLIMIIATFCEWMYVPRAWAGAQHLTRRLMFLIGMFALNVGPSVYIFGFSDQTGKIALA
LGIVQFFVALATFIFFSIMPLGGLFGSYLTSKKSRQYVASQTFTASWPRLSGNDMWMSYGLWVLVFAA
KMTESYFFLTLSLKDPIRILSVMEMRNCVGDKIVGTILCKYQPIVLLVLMFCTDLILFFLDTYLWYII
WNSVFSVARSFYLGVSIWTPWRNIFSRLPKRIYSKVLATTDMEIKYKPKVLISQIWNAIVISMYREHL
LAIDHVQKLLYHQVPSEQEGKRTLRAPTFFVSQEDHSFKTEFFPSQSEAERRISFFAQSLSTPIPEPL
PVDNMPTFTVMIPHYGEKILLSLREIIREDEPYSRVTLLEYLKQLHPHEWDCFVKDTKILADETSQFN
GENEKNEKDTARSKIDDLPFYCIGFKSAAPEYTLRTRIWASLRSQTLYRTISGFMNYSRAIKLLYRVE
NPEVVQMFGGNSDKLERELERMARRKYKIVVSMQRYAKFTKEERENTEFLLRAYPDLQIAYLDEEPPV
EEGDEPRLYSALIDGHSEIMENGMRRPKFR**IQLSGNPILGDGK**SDNQNHAIIFYRGEYIQLIDANQDN
YLEECLKIRSVLAEFEEMTVENVSPYTPGLPPPSTTPVAILGAREYIFSENIGILGDVAAGKEQTFGT
LFARTLAQIGGKLHYGHPDFLNGIFMTTRGGVSKAQKGLHLNEDIYAGMNALLRGGRIKHCEYYQCGK
GRDLGFGSILNFTTKIGTGMGEQMLSREYYYLGTQLPLDRFLSFFYAHPGFHINNLFIILSVQLFMVV
LINLGALKHETITCTFNKNLPITDPLKPTGCANLVPIENWVARCIVSIFIVFFISFIPLVVQELTERG

FWRAATRLAKHFSSLSPMFEVFVCQIYANSISANLSFGGARYIGTGRGFATARIPFGILYSRFAGPSI
YVGARLLMMLLFATMTAWGAWLIYFWVSLLALCICPFLFNPHQFAWNDFFIDYREYLRWLSRGNTRSH
SASWIGFCRLTRTKLTGYKRKALGDPSSKLSGDIPRARFTNIFFSEIISPLILVAVTLIPYLFINSQR
GVTADLNPDKTVEATNSLIRVGLVALAPIGVNAGVLAAFFGMACCMGPLLSMCCKKFGAVLAAIAHAI
AVIMLFAFFEVMFFLEGFSFAKALLGMITVLAIQRWIYKLIIGLALTREFKTDTANVAWWTGKWYSMG
WHSVSQPGREFLCKITELGYFAGDFILGHILLFIMLPALLVPMIDTVHSVMLFWLRPSRQIRPPIYSL
KQSKLRKRRVIRFAILYFVMLVLFIALIVGPLVAGKYITGFTLPLQLAQPTGLNRNDTLSSETGTAVN
GGAAATDAASSTVAARRLLARHY

>g16315: CBM43+GH72
MRGLSAVAGVAALFARSVVADLDPIVVKGAKFFYKTNGTQFFIQGVAYQQDYSTNGSSSTTASSAYTD
PLANAARCRVDIPLMKQLNMNTIRVYAVDPTQDHTACMQLLQDNGIYVVADLGQPGLSINR**DSPAWNT**
**QLYAR**YTSVVDMFAPYSNVIGFFAGNEVSNNKTNTNASAFVKAAVRDTKAYIKAKNYRQMYVGYATND
DAEIRANLETYFNCGDTSEAIDFWGYNIYSWCGDSSFTESGYDQRVAEFKNYSVPTFFAEYGCNTVQP
RKFSEVKAIYSSQMTGTFSGGIVYMYFQEANDYGLVQVSGSTVSQLADFTYLSSQMATIAPTGVQMAS
YTPTNSPQACPAVQTGVWEAKASPLPPVANAQLCSCMVNSLQCVVKSSQAENTYGTLFNQVCGYGSSC
AGIAAIASNATYGAYAACNSTQQLSFAFNQYYLSQNKASDACNFGGSATTKAASTPSGCASLISQAGS
AGTGTVSSGASSTATKKSAAGVTSVPSFNINLFGLGIYVSMAVVVGAGMILL

>g1789: GH17
MHAAQSFVALATLASVASAQVMGFNSGATLDTYKVKTQSDYEAEFTTAQGLVGAPGKFNSVRLYTMIQ
GGTDTDPTSAFQAAIKTNTTMLLGIWCSGTTTIEKELKALSTAITTYGAKFTDLVVGISVGSEDLYRT
SVTGIINK**SGIGNSPDAIVNFIK**DTRKAIANTPLSGTPVGHVDTWTDWTNSSNKAVIDAVDFIGNDLY
PYYEDTKDNSANNAVELFNEAYNATLAAAGGKPVWITETGWPTSGPLFGKATASNSDAQKYWQTVGCQ
LFGKTNVWWYNLRDSNPANEAKFAITSDLSTTPSFNLTCPAVVKTTKGGDGISSKSNSTSTATFPGST
GTSSSSSGNVTTGSSAGARTSGAAGSGTSPSSPVVTGVASSAGAMMGLTMCILLGSISLLL

>g21360: GH72
MKASSAFVATCALFSSVIAGSVSRR**ANTISNSNTPPVSVKGNAFFTSK**GRFYIRGVDYQPGGSSLLKD
PIADLDGCKRDVAKFKELAINTIRVYSVDNSADHDACMKLLADAGIYLALDVNTPYYSLNRKDNASIA
MSYNAVYLQSIFATIEAFAKYDNTLLFYSANEVINDDSTTFAAPYIKAVTRDMKAYMKARSLR**AVPVG**
**YSAADIESNR**YQTATYLNCGPDAARSDFFAFNDYSWCDPSSYTISGWDKKVATYSNYSLPLFLSEYGC
NKNKREFEEVKALYGSNMTPVYSGGLVYEYSQEEADYGLVDISGNTVTERPDFTALK**SAFAGTANPTG**
**DGGYK**SSGSASPCPAK**SDIWEVEDPTILPNIPSGATKYMTSGAGAGPGNK**GSTGSQTAGGASTGWSTT
SSSGSTTSGSASSATSSKAAAGNLQVPQLSMAPFVVAAVAGLSGLIGGAGFLL

>g21591

MLALHRTLLGLLAFASTSLAVNTVTIQGQDFVDTVTKNRVMIIGVDYQPGGQGGYDPNVRADALSNGT
VCLRDAALLQKLGVNTIRVYNVDPNANHDLCASIFNTAGIYMIIDVNSPQQSINRADPSSSYTVDYLT
R**IFAVVEAFK**GYPNTMAFFSANEVMNDIDTGKSNPPYIRAVQRDLRQYIAKNSQRTIAVGYSAADVRP
ILQDTWAYLQCNANSTDDWSRSEFFGLNSYSWCGADATYQTAGYDQLVSMFQNSSVPVFFSEYGCNKP
AGLARPFNEVQALYGPQMTSLSGGLVYEYSQEESDYGLVVINANGSITLRGDFDNLQNQYNKLNVTLL
QSTAAGNTQITPPQCSASLITNSGFSKDFTVPAQPSGAAALINSGISSPNQGKLISTGDLNVKQQIYS
SSGKLITGVAVKAVSGANTPGGENTSGSSATSTSTSSGTASPSASKKAAAASLQVTEGVMMRGLILAS
AIALGSGLIWRP

>g6964: GH3+CBM1

MKLSLVAAASLLVVSATATSPKLRSRQYSNSTTSSNPGAQFGQTSPPYYPSPWMDGSGGWETAYQKAQ
AFVKQLTLLEKVNLTTGVGWEGEACVGNVGEIPR**LNFPALCMQDSPLGVRSADYVSAFPAGGTVASSW**
**DRQVWYQR**GHDMGSEHR**DKGVDVQLGPVVGPLGR**APEGGR**NWEGFSPDPVLSGIAVAQTIK**GIQDAGV
IACTK**HFIGNEQEHFR**QSPEAASFGVNISESISANIDDTTLHELYLWPFADAVRAGTGSIMCSYNQVN
NSYACQNSYLLNNILKGELGFQGFVMSDWQAQHGGVSTSLAGLDMSMPGDTVFNSGISFWGANLTLAV
LNGTVPEWRIDDMATR**IMAAYYLVGR**DTK**KVPVNFASWTK**DTFAYRHPAVNSR**YELVNQHVDVR**AEHF
RNIRDHAAKSTVLLK**NSGVLPLTGK**EKFTGVFGEDADTSAWGPNGTPDRGSDNGTLAMGWGSGTADFP
YLVSPLTAIQNELVKNNALVQSVTDNWAYAQIASLASQVSTAIVFVNADSGEGYIAVDGNIGDRNNLT
VWRNGDTVIQNVTAKCNNTIVVIHSVGPVIVTDWYNNPNVTAILYAGLPGEQSGNSLTDILYGRYNPG
GK**LPFTLGAK**REDFGTDLLYTPNNGGNAPQDQFTEGVFIDYRHFDKAGIKPIYEFGFGLSYTTFAYSG
LQVQSHSVNTYTPTTGSTSAAPVLGTFSNNTADYLFPSNFTRVGLYIYPYLNSSNPATASQDKDFGKD
NLPAGSRDGSPQPRIAAGGGPGGNPQLYDVLFTVSATIQNTGQVEGDEVVQVYVSLGGPKDPVRVLRQ
FDR**LTIAPGATATFQADLTR**RDVSNWDTVAQNWVISNYTK**TVFVGSSSRTLPLSQTLTFGSY**

>g7121: GH17

MFAKLLALAALPAASNAFGVLK**GFNYGSTDASGVVK**DQAR**FEQEFSTAQNLVGTSGFNSAR**LYTTIQG
GTTNSPISAIPAAISTQTSLLLGIWTSAGQAIVDNEIAALKAAISQYGTAFTDLIVAVSVGSEDLYR**N**
**SGYPGASDPGPGANPDVLANYIGQVKAAIAGTSAEGR**LVGHVDTWTAFVNSSNNALISAADFLGVDAY
PYYESANGNDISNAANLFASAYSQVVAVAQGKPVWVTEAGWPVSGPTVAQAVASPENARSFWTSVGCN
QLFDKINVWWFQLDDYPTSPNPAFGVIGTAFSTTPLFDLSCPATTKSRRRSNRAA

### Glycoprotein and mannose metabolism

>g12843: GT66

MDALFQGDAAKNTRTLLRAIILLTIAGAAISSRLFSVIRFESIIHEFDPWFNFRATKYLVQHGFEPFW
NWFDDRTWHPLGRVTGGTLYPGLMVTSGVIYHFLRLISLPVDIRNICVLLAPAFSGLTAYATYLLTSE
MSTSPSAGLLAAAFMGITPGYISRSVAGSYDNEAIAIFLLVYTFYLWIKAVKEGSVMWGALAALFYGY

MVSAWGGYVFITNLLPLHAFVLICMGRYSARLYVSYTTWYAIGTLASMQIPFVGFLPIRSSEHMSALG
VFGLLQIVGFVEYVRLQLPSKQFQTLLRSLVLLIFLVSFGGLVLLTVSGVIAPWTGRFYSLWDTGYAK
IHIPIIASVSEHQPTAWPAFFFDLNLLIWLFPAGVYLCFRTLKDEHVFIVIYAVLSSYFAGVMVRLML
TLTPVVCVAAAMALSQILDIYLLAESPSEELQTLSSAEAAK**AAAGTSLLSDGLR**STTKPIVGIYSYMS
KATVVVCSTIYLLIFVAHCTWVTSNAYSSPSVVLASKMPDGSQHIIDDYREAYYWLRQNTPQNAKVMA
WWDYGYQIGGMADRPTLVDNNTWNNTHIATVGKAMSSREEVSYPIMRQHEVDYVLVVFGGLIGYSGDD
INKFLWMVRIAEGIWPDEVKERDFFTPRGEYRVDDEATPTMKNSLMYKMSYYNFNALFPAGQAQDRVR
GSKLPAQGPELSTIEEAFTSENWIIRIYKVKDLDNFGRDHSNAVAFEKGHKKKKAAKRRGPRSLRLE

>g18706: GH31

MLQMGTQSKGWSRTLSLLCLVGLFTPVFTVKHENFKTCDQSGFCKRNRQYADAAGTAAFTSPYELESS
SISFQNGQLKAAVIKTVGKSGEKVRLPVTISFLESGSARVTLDEEKRVKGDIELRHNSKARKERYNEA
GQWAIVGGLAPSAGAALNNAAEKGTTIVKYGPSGTFEALVRHAPFSIDFKRDGETQIQFNEGGLLNME
HWRPKIEKKVEEPKEGEAGPAAPAPEDPNAEDEGTWWEESFGGNTDTKPRGPESVGLDISFPGFEHVY
GIPEHASSMALKQTRGGDGAYSEPYRLYNADVFEYELDSPMTLYGAIPFMQAHRKGSSVGVFWLNGAE
TWVDVVKSKTNANPLSLGIKGSTTTQTHWYSESGQLDVFVFLGPTPKDVIKSYGELTGYTQLPQEFAI
AYHQCRWNYVTDDDVIDVDKKFDKFKIPYDVIWLDIEYTDGKKYFTWDPLTFDDPENMGKQLAKRERK
LVTIIDPHIKNTDSYHVVDQLKSKGLAVKNKDGDIYEGWCWPGSSHWVDCFNPAAIAWWKSLFAYDKF
RGTLPNTFIWNDMNEPSVFNGPETTMPKDNLHHDNWEHRDVHNINGMTFQNATYHAMLARNKAEEKSP
RRPFVLTRSFYSGSQRVGAMWTGDNLAEWSHLAVSLPMILNQGISGFPFAGADVGGFFGNPSKELLTR
WYQAGAFYPFFRGHAHIDTRRREPYLAGEPYTSIITKALQLRYALLPSWYTAFHEASTTGAPIVRPNF
YVNPADEAGFTLDDQLYLADTGLLFKPVVTEGAESVDIYLGDDAPYYDYFDYTIVKGKGSHTLKAPLD
KIPLLMRAGHIFPRRDRLRRSSGLMKLDPYTLVLVLGPDGKAEGELYVDDGESFDYEQGAYIHRKFIY
ENGSLRSEELGKKGKLTDKYAKKMEKVRVERVVIVGAPSAWKGKKQVLVSEEGEKDSKGGKKVK**IDFT
DATAGK**AAFAVVRDPKVAVGKGWKIDFGA

> g2667: GH63

MLTPKFASMISGLLLVSTAIATPLSSPGNHQKRAIPRSSVPPTMAPYDRADYVYDPEKEESGFDRRTA
RTFDSDAGASHTTKRSPTDLERRYDGWEATPICNKESFGGKADYNQVNELMDTLLMQNGKPEVGAGPK
KCAMAGSTLGLSLSLTFGLRASRSFISRTNTTIANAIMRSRLQSPSPWPSLLSSLLLLLIAPITSAST
SQPTTIADIERASNQSLLWGPYRPNLYFGVRPR**IPESLLIGLLWAK**VEDYQSVQHNFRHTCEQHEGMA
GYGWERYDPRHGGVQTIHDAGNQIDITTSFFKETGEGSGDRGGNWGVRIKGVPRADAQEDLKSTVVFY
ASTEGQGLNNRLEVKNVDELQGGGFDGDVVLKGENLGLGEYKIVVKGDEGTENKHPSVTHPSGSEKDL
GKTLVKSSTVPEDAIWQTKPILFAMLKEQIDEYVEKYTKENPPPPFQLYTIKSEAGSGNVHFVQKVFE
GAFEFDVLFSSSAAESELASSDLSKGISEIVKTFDTRFDSIFKPAAPFNSGKYLDFSKSLFSNLLGGI
GYFHGNSRVDRSYAPEYEEDNEGFWEEAAEARARADVK**LEGPNELFTSIPSRPFFPR**GFLWDEGFHLM
PVVDWDIDLTLQIVK**SWFALIDEDGWIGR**EQILGAEARSKVPEEFQVQYPHYANPPTLFFILSAFVDK

LTETAPSSKDAEYSPQLLDKEVATSYLKELYPLLKRHYNWFRKTQQGDISSYDRKAVNTKEGYRWRGR
TPRHILTSGLDDYPRAQPPHPGELHVDAISWVGLMATSLQKIGLFLNEKEDVEKYTKQLTGIRSNIAD
LHWSEKDGVFCDATIDEFEENSLVCHKGYISLFPFMLGLLDPADDGNKIAKILATIGNKEELWSEHGI
RSLSIADEAYGTDENYWRSPVWINMNYLIVSR**LVALAQDPTAGSDNQK**TATKLYTDLRINLVETVYKS
WKETGFAWEQYNPETGAGQRTQHFTGWTSMIVKILGMPDLSKGSAKVRDEL

>g3844

MGGSHDTPTSTKYPTLVQKPVGKQIHNLYLDRLQQFTDNGQYRKQGLLDKIIEARASGDQWVRLEVYS
PPDLSRPTFKEATSHKFRDTHVGESFGPSWATHWFKIHLTIPDDLAKKEHLEFVWDANNEGMIWTEKG
DVVHGLTGGGDR**TQWILPESWR**DGKEHIFYVEMACNGMFGNAPGGDSIQPPPPDKYFQLHTAEITAVN
LDARQLYIDFWIIGDGAREFPGDGWESHKALQVCNAIMDCFIAGQGTKECIKECRKIAREYIGNVDTP
KIYDGDLPSLVTAVGHCHIDTCWLWPWAETKRKVARSWSNQCDLLDRYPELRFCASQAQQYKWLEMLY
PSLFDRVKEHVKKGNFQPIGGSWVEHDTNMPSGESLVRQFVYGQRFFESHFGQRCTTFWLPDTFGYSS
QLPQLCRLAGMSRFFTQKLSWNNINNFPHTTFNWVALDGSQVLCHMAPSETYTAEAHFGDVKRSITQH
K**SLDQDETSLLVFGK**GDGGGGPQWEHIEKLRRARGISDTVGLLPRVK**LGDSVDDFFAK**LEKKAETGTD
FVTWYGELYFELHRGTYTTQANNKRNNRKSEFMLRDIEFLATMATIKDDVDGKKSTYKYPKKEIDFMW
EAVLLCQFHDCLPGSSIEMCYDDSDKLYAEVFETGTKVLTDALSELGFDDDKKTSSVGDLVALNTLPW
ARSEMTRLPIKSEAPKYAAIDSTHTGLGVVRALTAASSAPVSIRETEK***GSFELSNSAFDVK***MSDGVIT
SLFDKRANREVIAKGGKANQLVIFDDKPLYWQAWDVEVFHLQSRQELSSSTSKIAEQGPHRVSVVTET
KISAESWVKTTISLNAATDDNYASIDVEAEVEWHETMKFLKVEFPVDVSNTEASYETQFGIVRRPTHY
NTSWDMAKFEVCCHKWADLSESNYGVSILNDSKYGFATCGNLMRLSLLRAPKAPDGHADMGRHQIRYS
IFPHNGPLDYKTIRAGYSFNNPMKLHHHPKPASISSLLSSFNIDGSKSLIIDTVKRGDDDVDVSRGEL
PAKKGKSVILRIFDALGGKSK**GILTWGDVPVK**AVFK**TNLLEDAGDELVVLK**SGKGVEIELRAFEVATF
RLELQ

>g9548: GT24

MRVPSWLLPAEILLLVGSLGLVPLTGAAPSINVGLHASFNSAPYLVELLETAAEEKPDVYYKILDRIS
DGYFSDASTDKELYEKFLRLLKYDNLITDSESLSSFELALSIHNAAPRIEAHYQYYKTTIEPLLEKSK
GKDCETWLAFTGKQFCSPQFEKADATIK**GINIEGGVPILPFDR**IFGSPTTIPAVIYADISSPVFKKFH
SLISEAAKNGRVSYRVRHKPSKSERKPLEMSGFGTELALKRTDYIVIDDRKAEEGKEADKESSKPADI
DLVDEEVADLKPLTSSEVTELGLK**ASTYVLGSEDPLETLVK**LTQDFPKYSSVIAGVNSSEAFLEEHRE
NRALLLPAGFNVIWINGVQFDSRKVDAFSLLDHMRRERALLGSFQEMELTSEEAIKLLTHPSIAEAQS
DVDVQRYDWRDETEGGNVIIWMNDIEKDKRYAAWPSEIYGLLQRGYPGQLPTVRKDIHNAIIPIDFSE
PTSLHRATETVQDFVKRKIPIR**FGLVPITSTAGAAK**QAQVVYHILDTYGLGALMQYLESSLSAKKTAA
PHESTFKAVIEKRKPKAGREAVALQAVLQSEESAIQVGGAKKYLTRLGATGSTPPVFMNGVPIKNDDE
WLQAMVQR**VSQDLQWLQR**GVFEESITQDMWIPSNWLNDSSIRRNALVIPENHKDIK**LLNLLNVLK**WED
EQTMKIMPTIPADEDSDKAKWAQLVLVGDFDTQVGLQMILDAIKFRRENPNIELVLVAQSGSPMIKSP

RIVQIWPKETKWTMETIQAVYRDIKALLAAPYDHAAHDNSQFWPVETEQLAQAFGFDQGQNGLIINGR
KIGPISAESAFTKDDFAALYKFEFKKRIAPASEAIAELEYKDKIKTVSDAAKLCSLLAVSTVSDMPEG
IFELPPPLRTRAYYEWSDSETAVTVGDNTTALINIVAVLDPAAEPAQRWSPILRVLSKLEGVSLKLFL
NPKENLHELPVKRFYRYALNEKPSFDDDGAVTAPGVKFEGIPKDTLLTVAMDVPPSWLVAQK**ESLYDL**
**DNVKLSSLPLGDNVDATYELENILIEGHTR**DSKGGDQPPRGAQLVLSTAKDPHFADTIIMANLGYFQF
KANPGYYNITLKPGLSSKIFKLDSAGALGYEATAGDEISEVCLLSFQGLTLYPRFSR**NPGMEDEDVLG**
**STKSSTASELASK**GADLVDGFLNKAGIKKTKGVQSAQADINIFSVASGHLYERMLNIMMVSVMRHTQH
TVKFWFIEQFLSSSFKSFLPTLAKEYGFKYEMVTYKWPHWLRAQKEKQREIWGYKILFLDVLFPLDLD
K**VIFVDADQIVR**TDMYELVTHDLKGAPYGFTPMCDSRVEMEGFRFWKQGYWKNFLRGLPYHISALYVV
DLKRFRQIAAGDRLRQQYHQLSADPASLSNLDQDLPNNMQMMLPIHSLPQEWLWCETWCSDESLKDAK
TIDLCNNPQTKEPKLDRARRQVPEWTEYDDEIAALARRTKQSEKVPTGDEQAGTEGRPKESVVESESA
ESTHVRDEL

### N-acetylglucosamine metabolism

>g21234: CBM18+GH18

MKLSTTLSLLTAGLAAASSCTKRTTTGRNVMYYDQYHTNLTTLTPEIASGITHVIIAFIPSTNFTVAN
TSAFVPFESVSKVRTRFGNSTKVMIAIGGWGDTAGFSIGAKTNESRALFAKNVKKMLLETGANGVDLD
WEYPAGNGADYKTNPNSNKTDEIITYPLLVQAISEAIGSSYLLSAAVPGLAR**DMIAYTTATGPAIFK**Y
LDFVNLMSYDLMNRRDTVMKHHTSIADSITAVDLYTSIGLAPSKINLGIAFYAKWFAAAPNGTCDAAS
PLGCKTALLEDPVTGDDLGLAGAVTFEASNYAIVDETKLANSTDGSCGANVGPFGTRCIPGNLDWISR
ESNPVQSSGITDLGRTIALGRQLVFQSDLNKKWPGASCVTLVPPESNSSSTLAASQEILPGDTTRMEF
QEAAAAFRYRS

>g2198: ViCHS3b

MAHQGYGGGGYNDPSQPPGSQYHAPGSRRGSDEEHEVAQSLLHADPTGTREGPFGNQYASEHTDRMRT
PEVRPTSTYSLSETYADNTGYGPGYNQGYAEDQQAGYDMPPRIASPYSRSETSSTEAWRQRQQPGGGA
AAGGGLKRNATRKIKLAQGAVLSADYPVPSAIQNAIQAKYRNDLEAGSEEFTHMRYTAATCDPNDFTL
KNGYNLRPAMYNRHTELLIAITAYNEDKVLTARTLHGVMQNIREIVNLKKSEFWNKGGPAWQKIVVCL
VFDGIDPCDKGTLDLLATVGVYQDGLMKKDIDGKETTAHIFEYTTQLSVTANQQLIRPLDDGATTLPP
VQMMFCLKQKNTKKINSHRWLFTAFGRILNPEVCILLDAGTKPGPK**SLLALWEGFYNDK**NLGGACGEI
HAMLGRGWK**NLLNPLVAGQNFEYK**ISNILDKPLESSFGYVSVLPGAFSAYRYRAIMGRPLEQYFHGDH
TLAKILGPKGIDGMGIFKKNMFLAEDRILCFELVAKAGSKWHLTYVKASKGETDVPESAPEFISQRRR
WLNGSFAASIYSLMHFGRFFKSGHNPIRKFFFVLQMFYNVAMLVLSWFMLGSFWLTTSVIMDLVGGTV
DEIKNPTATTASKGWPFGIKYSPHVNAVLKYIYLGFVILQFILALGNRPKGSRISYIVSFCVFAIIQL
YLIVLSFYLLGKALSGNTVKDNFNDLEKFFSPDGVGVILIAVIATFGLYYIASFLYFDPWHMFTSFPQ
YLLLMPSFTNILNVYAFSNWHDVSWGTKGSDKSEALPSAKTEKSSDGKHTVIEEVDLAQADIDSQFEA

TVKR**ALSPFVATPEDNTK**TTDDGYKSFRTKLVSTWIFSNIIIVIVITSETFDFIFPASSASTKRTATF
FTALLWVTAGLSVIRFSGCIFFLFKTGALRITRKR

### Trehalose metabolism

>g17565: GT20

MSLEPLQMEGRLLLVSNRLPITIKRNDEGKYDFNMSSGGLVSGLSGLKNDVTFEWYGWPGLEIPDDEV
GDLKTKLKEEYNAVPIMLDDELADRHYNGFSNSILWPLFHYHPGEITFDESAWEAYTEANRLFAKAIA
KDVQDNDLVWVHDYHLMLLPAMLREELGDTKKNVKIGFFLHTPFPSSEIYRILPVRNEILLGVLHCDL
IGFHTYDYARHFLSSCSRILGLPTTPNGVEYKNK**VVTVGAFPIGIDPEK**FAEGLKKPKVIERIETLKR
KFQGVKLIVGVDRLDYIKGVPQKLHALEVFLTEHPDWIGRVVLVQVAVPSRGDVEEYQNLRSVVNELV
GRINGKFGTIEFMPIHFMHKSVSFDELVALYAVSDVCLVSSTRDGMNLVSYEYIATQAERHGVMVLSE
FTGAAQSLNGSLIVNPWNTEEMAEALHEAVTMGDEQRKINYDKLARYVNKYTSSWWGQTFVTEMVRMT
EQTEKKLSIRSGSKVAFADKESAASTEEKANEPTGGLTVTDGHLVSSSSASDSPTSGGEANEKPVVTS
PESPVKEAPETEGNRAPAESATPEKSHTL

>g315: GT20

MTSPPKGKQGDAQETGLLSRTRTNSDDSFHAHLVSNAVPVTPGVHTAGHSAYFEQKREEGEPDQEEFH
DDDASPGPNWNANNYHVPKTDDGPPVSPGLAATDAKTAQEAIRKLTMAASGDSGKKELSDVDPRAAHP
QLGLSGHIISATFVVPYNISFAPGKDWELKPRSGTSALFDSFSYLASSSSPWNHTLLGWTGEIKKNPA
AFPSNPALAAMENLNSTKKTSIPVEGKMKLVDPTQSTSMKVSRKDRQRLETQLERDHGGKIVPVWLVD
DVSEDDDIYTIENQSRWRSFAERELYTLFHYKQNEPSDGRGVRTAWADYYKLNKLFADRIIEVYKPGD
IIMIHDYNLMLLPSLIRQRLPKAYIGFFLHIPFPSSEYYRCLSRRKEILEGVLGANLIGFQSFTYSRH
FSSCCTRILGFDSSSSGVDAYGVHVAVDEFPIGINAFSTKKAAYNDPMVEEKMAGILQLYAGKRIIIG
RDRLDTVRGVSQK**LEAFENFLER**YEEWRDKVVLIQITSPNATNTVDDGGESKFMDKISDRVSKINGKY
GSLSFSPVKHFPQYLSKEEYFALLRVANVALITSVRDGMNTTSMEYVVCQENNFGPLILSEFSGTSGS
LKAAIQVNPWDLGGVADALNSALHMSNEERKEKHAKLYRHVVNNNVQNWTNKLFKRFLINLESFDQSF
STPALDRVKLLTAYRTAKKRLFMFDYDGTLTPIVKNPESAIPSDRVLRTLK**TLAADPANTVWIVSGR**D
QAFLDHYMGHISALGLSAEHGCFMRQPESDDWENIAAHMDMSWQQEVKNTFNTYTDKTPGSHVEVKKV
ALTWHYRNSHPELGLEMSRKCQRELETTVARNHDVEVMTGKANLEVRPKFVNKGEIARRLVKAYGSGP
GDAPEFVFCLGDDSTDEDMFRALKQSELPSGNVFSVTVGASSKQTLASWHLLEPHDVISCISLLNGSI
DNENVGAVSVVDGTIPEMTEARI

### Galactose metabolism

>g20534: CBM87+CE18

MFALRRLGSALVASLSLASLVSAANTVNSTILVFARDTASGYSGTSGLAGYGIPYQLVVVPQAGITLP
VLNSTATAGNFGGIIILSDVAYSYDSGWASAITAAQWAQLFAYQTSFGVRMVRLDVYPGSNFGTTTAI

AGEGCCDAGVEQLVSISSNTAFPKAGMNTGAGVTTQGLWHYPAIINNASIATEIAQFAPAGDFTTTTT
AAVINNIGGRQQMAFFMGFATDWSSTSNFLQHAYIHWMTRGLFVGRRRIYFNTQIDDMHLVTDIYQPA
GSLYRVVPNDMVTHVSWVNGLNSR**LPAGSSYK**VEIGHNGNGAIENALTIDPTSCTPNSAIEYAEQIDT
PLEFQKPLGTGTNIWPATPTLYPAGWGTACLNKDPLAAWFRVAANRNAFFHISHTFTHEGLNNATNSD
ANKEIAFNRAWFYYMGLDSAATFSTTGIIPPAITGLHNGDVIRAWIANSIMHVVGDNTRPPLLNTVNE
FWPLTSTVAANGYAGLTILPRWATTIFFNCDTAACTTAEWVNTSGGKGDFAALLANAKATNTRHLFGL
HQDGFMFHQANLRADSAIPSYTVGSQSVQSLLQIWVETITQEMSRLTTWPLISLRQDDMATQFRNRQT
RDGCSPNMVWNYSADNKKIVGATVTANRNTCSVPIFATFPSALTTSPGTNDGVGSDSLQYPITLSGAA
KTYTFSTAISV

### Effector candidate

>g18338: ViEcp6 LysM

MLFAKSSVAMVSLFTLLVAAAPAPELLSARELCNGTITLDERVQK**YTIASGDSLGAIATK**FNRGICDI
ATANKITNINFVTAGQVLTIPAQVCIPDNASCQPKTPEATATSILGGPGFYIVVSGDTLTAIAKNFQI
TLQSLIAANPAITNPDLILVGQVIIIPVLPGSSCTISPYVIKSGDIFFDLAAK**FGTTAGQLLSLNTGT**
**DPTK**LAIGQTVTIASGCKNATATTGENKYWKGQWGGPGGKWEGKWGN
